# Acute, chronic and conditioned effects of intranasal oxytocin in the mu opioid receptor knockout mouse model of autism: social context matters

**DOI:** 10.1101/2024.03.07.583912

**Authors:** Fani Pantouli, Camille Pujol, Cécile Derieux, Mathieu Fonteneau, Lucie P Pellissier, Claire Marsol, Julie Karpenko, Dominique Bonnet, Marcel Hibert, Alexis Bailey, Julie Le Merrer, Jerome AJ Becker

## Abstract

Autism Spectrum Disorders (ASD) are neurodevelopmental disorders whose diagnosis relies on deficient social interaction and communication together with repetitive behaviours. Multiple studies have highlighted the potential of oxytocin (OT) to ameliorate behavioural abnormalities in animal models and subjects with ASD. Clinical trials, however, yielded disappointing results. Our study aimed at challenging hypotheses accounting for such negative results by assessing the behavioural effects of different regimens of OT administration in the *Oprm1* null mouse model of ASD. We assessed the effects of intranasal OT injected once at different doses and time points following administration, or chronically, on ASD-related behaviours in *Oprm1^+/+^and Oprm1^-/-^* mice. We then tested whether pairing intranasal OT injection with social experience would influence its outcome on ASD-like core symptoms, and measured gene expression in several regions of the reward/social circuit. Acute intranasal OT improved social behaviour in *Oprm1^-/-^* mice at a moderate dose (0.3 IU) shortly after administration (5 min). Effects on non-social behaviours were limited. Chronic OT at this dose maintained beneficial effects in *Oprm1* null mice but was deleterious in wild-type mice. Finally, improvements in the social behaviour of *Oprm1^-/-^* mice were greater and longer lasting when OT was administered in a social context, while the expression of OT and vasopressin receptor genes, as well as marker genes of striatal projection neurons, was suppressed. Our results highlight the importance of considering dosage and social context when evaluating the effects of OT treatment in ASD.

**Highlights:**

- Acute intranasal oxytocin improved social behaviour shortly after administration
- Effects of oxytocin on non-social behaviours were limited
- Chronic oxytocin maintained beneficial effects in mutant mice but was deleterious in wild-type mice
- Oxytocin triggers greater and longer-lasting improvements in social behaviour when administered within a social context
- Chronic oxytocin suppresses the expression of its own receptor gene and key striatal neuron gene markers

## 1 INTRODUCTION

Autism Spectrum Disorders (ASD) are highly heritable neurodevelopmental disorders characterized by impaired social communication and interaction associated with a restricted, repetitive repertoire of behaviours, interests and activities [1]. Alongside these core symptoms, ASD is frequently associated with comorbid symptoms such as high anxiety, cognitive impairment, motor stereotypy, aggressive behaviour, abnormalities in pain sensitivity and epilepsy [2, 3]. Despite the identification of vulnerability genes and environmental risk factors [4, 5], the etiopathological mechanisms underlying ASD remain essentially unknown. To date, approved pharmacological treatments for ASD mostly target associated symptoms [6, 7] and evidence-based behavioural interventions remain the only treatments proved to ameliorate core social deficits [8].

Among potential pharmacological treatments for ASD, oxytocin (OT) stands out as a highly promising molecule to relieve socio-communicational impairments. OT is a neuropeptide synthesized in the paraventricular (PVN) and supraoptic nuclei (SON) of the hypothalamus. Animal research has evidenced, beyond its key contribution to reproductive functions, a crucial role for this nonapeptide in shaping social behaviours, including social approach and reward [9, 10], social recognition [11] and memory [12], parental behaviour [13], pair bonding [14] and emotion discrimination [15]. Consistent with this, targeted disruption of genes encoding OT and its receptor (OTR) impairs social behaviour in mice [16, 17]. Experiments in mice have shown that OT is released in response to social cues, making it a key contributor to the rewarding properties of social interaction [6, 9]. In human studies, OT effects have been examined after intranasal administration as oral intake does not allow for sufficient bioavailability [18, 19]. In healthy subjects, intranasal OT increases social salience [20, 21], improves facial emotional recognition [22] and promotes in-group cooperation and trust [23, 24].

Beneficial effects on social behaviour under physiological conditions bode well for therapeutic effects of exogenously administered OT in ASD. Preclinical studies in mouse models have evidenced improvements in social interaction [25, 26], social preference [27-30] or social memory [31, 32] under OT treatment. Such prosocial effects persisted upon (sub)chronic administration or when OT was administered early in life [26, 27, 30]. Accordingly, in patients with ASD, acute intranasal OT application improved sustained eye gaze and social cooperation, but had limited effects on non-social behaviours [33, 34]. Unfortunately, clinical trials testing long-term, daily OT exposure in ASD yielded inconsistent results, with the largest to date clinical trial showing no effect of intranasal OT on social behaviours in children with the condition [35, 36].

Several factors may have contributed to these disappointing outcomes. Most clinical trials omitted to consider the heterogeneity of ASD, as chronic OT could be beneficial in a small subset of individuals [37, 38] and to test multiple or individualized doses [36, 39]. Moreover, animal studies suggest that the regimen of OT treatment needs to be considered, as chronic administration leads to severe social deficit in wild-type mice [40]. Finally, OT does not always behave as a facilitator of social behaviour [21, 41] and its prosocial effects would depend on social context [42], as predicted by the social salience hypothesis [21]. Taking all the above into consideration, it appears that chronic intranasal OT treatment, at a dose and social context not individually adapted, may not be optimal as an ASD therapeutic approach [36, 43, 44].

In this study, we challenged previous hypotheses by assessing the behavioural consequences of varying the dose, timing and context of intranasal OT administration in the *Oprm1* null (*Oprm1^-/-^*) mouse model of ASD. The *Oprm1^-/-^* is a well characterised mouse model of ASD mouse [45-48] with an altered oxytocinergic system [45, 46, 49, 50]. Upon acute administration of OT or a non-peptide analogue, these mice show rescued communication and social interaction [50, 51]. Here, we evaluated the effect of a range of doses of single intranasal OT administration on social behaviour in *Oprm1^-/-^*mice and their wild-type counterparts and assessed the contribution of OTR in mediating these effects using a novel selective non-peptide OT antagonist. We also tested whether acute OT would affect other, non-social, autism-sensitive behaviours. We then evaluated the effects of intranasal OT administration in a chronic setting. Finally, we assessed the behavioural consequences of pairing intranasal OT injection with congener versus object presentation in *Oprm1^-/-^*mice and *Oprm1^+/+^* controls. To gain insight into the molecular substrate of OT effects, we measured gene expression, notably for genes related to the oxytocin/vasopressin system, in several regions of the reward/social circuit.

## 2 METHODS

### 2.1 Ethics

This study was approved by the Comité d’Ethique pour l’Expérimentation Animale de l’ICS et de l’IGBMC (Com’Eth, 2012-033) and Comité d’Ethique en Expérimentation animale Val de Loire (C2EA-19). All experimental procedures were conducted in accordance with the European Communities Council Directive 2010/63/EU. Animal studies are reported in compliance with the ARRIVE guidelines [52] and with the recommendations made by the British Journal of Pharmacology [53].

### 2.2 Animals, housing conditions and breeding procedures

Equivalent numbers of male (25-32 g) and female (22-28 g) *Oprm1^+/+^*and *Oprm1^-/-^* mice [54] were bred in-house on an identical hybrid background: 50% 129SVPas - 50% C57BL/6J. *Oprm1^+/+^*and *Oprm1^-/-^* pups (F3) were bred from homozygous parents (F2), as we previously showed that parental care has no influence on behavioural phenotype in these animals (cross-fostering experiments [45]). Homozygous parents were bred from heterozygous animals (F1), to prevent genetic derivation. Mice in the same cage were of the same genotype: this breeding scheme likely exacerbates behavioural deficits in mutant animals by maintaining them together during early post-natal development [49]. We defined sample size (GPower 3.1) to ensure sufficient statistical power using ANOVA or Kruskal-Wallis analysis of variance to detect significant effect on our parameters (effect size f=1.80, α=0.05, σ=5, n=8, power=0.96). Cages of *Oprm1^+/+^* and *Oprm1^-/-^*mice were assigned randomly to a treatment group by the staff of the animal facility (blind to experimenters), provided that sex ratio was equivalent between groups, and that mice from different litters would meet during the direct social interaction test. Except otherwise stated, all mice were group-housed and maintained on a 12hr light/dark cycle (lights on at 7:00 AM) at controlled temperature (21±1°C); food and water were available *ad libitum*. Cardboard igloos (Dietex®, Argenteuil, France) and laying were provided in each cage as enrichment. Routine veterinary care and animals’ maintenance was provided by dedicated and trained personnel. At the end of experiments, mice were killed either by exposure to rising concentrations of CO_2_ over 5 min or cervical dislocation when brain samples were collected (qRT-PCR experiments).

### 2.3 Drugs

Mice received either vehicle (NaCl 0.9%) or oxytocin (PubChem ID: 439302; reference #03251, Sigma Aldrich, Saint Quentin, France) at the dose of 0.15 IU (400 µg/kg), 0.3 IU (800 µg/kg) or 0.6 IU (1600 µg/kg) via the nasal route, either acutely (5, 15 or 30 min before testing), chronically (once a day for 17 consecutive days, 5 min before testing) or repeatedly (6 administrations, 2-3 days apart, 5 min before testing). A solution of 1 IU contained 1.667 mg of synthetic OT. The doses tested in this study were similar to high doses of intranasal OT given to adolescent prairie voles [55] and in the lower range of the daily dosage used in clinical trials [35, 36, 56]. To assess whether the OT effects were mediated by OTR activation, we administered via intraperitoneal route either vehicle (carboxymethyl cellulose 1% in NaCl 0.9%) or the highly selective OT receptor antagonist LI183, synthesized in-house, at the dose of 7.5 or 15 mg/kg (synthesis and pharmacological profile in Supplement 1) 30 min before behavioural testing.

### 2.4 Intranasal oxytocin administration

For intranasal OT administration we adapted our protocol from (Huang et al., 2014). OT was dissolved in saline vehicle (0.9% NaCl) and administered intranasally in a volume of 5 μl to each mouse. Control mice received the same volume of vehicle (NaCl 0.9%). Dilutions were calculated so that 5 µL of solution contained either 0.15, 0.3 or 0.6 IU. We used Eppendorf tips (P10) to gently drop 2.5 µL of solution on each nostril. A 30 s handling ensured a reflex inhalation for each mouse.

### 2.5 Behavioural experiments

Equivalent numbers of naive male and female animals were used in each group. For acute experiments (Figures 1 and 2), each behavioural test was performed in an independent cohort of mice. For chronic experiments (Figures 3-5), behavioural tests were performed successively in the same cohort of mice, and testing order was chosen to minimize the incidence of anxiety generated by each test on later assays, except for evaluation of nociceptive thresholds. Indeed, the effects of chronic OT on nociception were assessed in a dedicated cohort of mice (see Figure 3a). Experiments were conducted and analysed blind to genotype and experimental condition.

**Figure 1.**
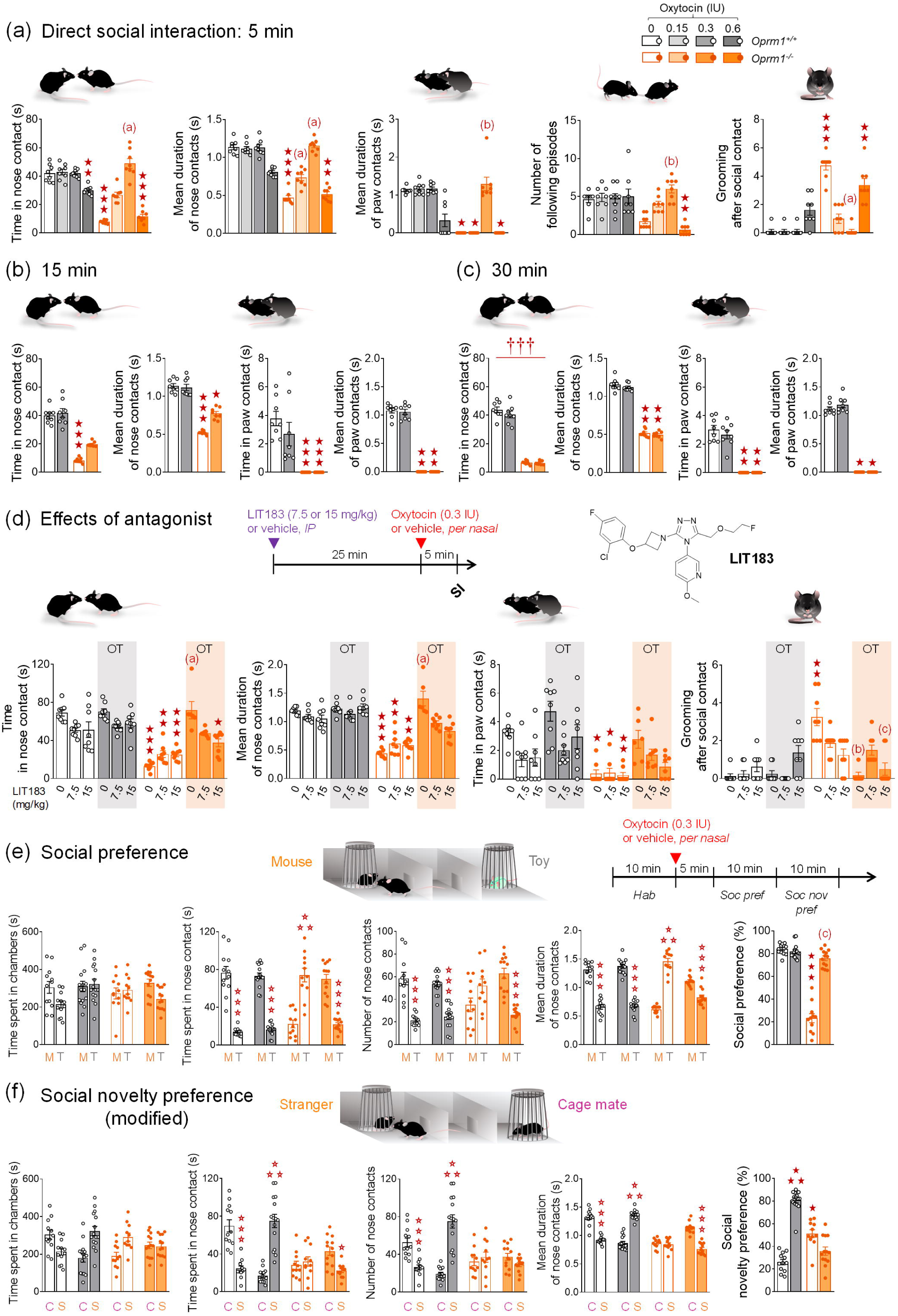
Acute per nasal administration of OT dose-dependently restored social behaviour in *Oprm1* null mice. (a) *Oprm1^+/+^* and *Oprm1^-/-^* mice received OT or vehicle (4 males – 4 females per genotype and treatment), via per nasal route, 5 min before the direct social interaction test and at the dose of 0, 0.15, 0.3 or 0.6 IU. In this test, vehicle-treated *Oprm1^-/-^* mice displayed a deficit in social interaction; OT at 0.3 IU fully restored the time spent in (*Genotype [G] x Dose [D] interaction: F_1,56_=39.80, p<0.0001*) and mean duration of nose contacts (*H_7,64_=53.13, p<0.0001*) of these mice, as well as their number of following episodes (*H_7,64_=37.57, p<0.0001*). It increased the time they spent in paw contact (*H_7,64_=52.71, p<0.0001*) and the mean duration of these contacts (*H_7,64_=49.93, p<0.0001*). A partial restoration was observed for the dose of 0.15 IU and beneficial effects were lost at 0.6 IU. Similarly, OT at 0.3 IU suppressed grooming episodes occurring after a social contact (a sign of social discomfort, *H_7,64_=49.76, p<0.0001*) while OT was only partially effective at 0.15 IU and had no effect at 0.6 IU on this parameter. (b) When administered 15 min before testing (4 males – 4 females per genotype and treatment), the optimal dose of 0.3 IU had only partial effects in alleviating social interaction deficit in *Oprm1^-/-^* mice, as evidenced by increased time spent in nose contact (*H_3,32_=28.28, p<0.0001*) and mean duration of nose contacts (*H_3,32_=26.28, p<0.0001*), but had no effect on the time spent in paw contacts (*H_3,32_=27.31, p<0.0001*) and their mean duration (*H_3,32_=26.59, p<0.0001*). (c) When administered 30 min before testing (4 males – 4 females per genotype and treatment), per nasal OT at 0.3 IU was ineffective in relieving social interaction deficit in *Oprm1* null mice. (d) The non-peptide OT antagonist LIT183 (see Supplement 1) or its vehicle (doses of 0, 7.5 or 15 mg/kg) were administered intraperitoneally 25 min before per nasal OT administration (0.3 IU) and 30 min before direct social interaction test (4 males – 4 females per genotype, LIT183 doses and OT treatment). Pre-test administration of LIT183 similarly tended to reduce the time spent in nose (*H_11,94_=71.65, p<0.0001*) and paw contacts (*H_11,94_=49.21, p<0.0001*) in vehicle and OT injected *Oprm1^+/+^* mice, without reaching significance. In *Oprm1^-/-^*mice, LIT183 blunted the effects of intranasal OT on the time spent in nose contact and mean duration of nose contacts (*H_11,94_=74.35, p<0.0001*), as well as on grooming after social contact (at a low dose of LIT183, *H_11,94_=57.01, p<0.0001*); paradoxically, however, OT antagonist reduced such grooming in mutant mice treated with vehicle. Of note, OT effects on the time spent in paw contact in *Oprm1^-/-^* mice did not reach significance in this experiment. (e) We performed a modified version of the 3-chamber test (*Oprm1^+/+^*vehicle: 6 males – 7 females, *Oprm1^+/+^* OT: 7 males - 8 females, *Oprm1^-/-^* vehicle: 5 males - 8 females, *Oprm1^-/-^*OT: 6 males – 8 females). During the social preference phase, intranasal OT increased the time spent in nose contact (*Genotype [G] x Dose [D] x Stimulus [S]: F_1,47_=76.4, p<0.0001*), the number (*G x D x S: F_1,47_=26.19, p<0.0001*) and the mean duration of nose contacts (*G x D x S: F_1,47_=124.01, p<0.0001*) with a stranger mouse versus a toy in *Oprm1* null mice, resulting in fully restored social preference ratio (*H_3,51_=30.35, p<0.0001*). No effect was detected in *Oprm1^+/+^* mice. (f) During the modified social novelty preference phase, vehicle-treated *Oprm1^+/+^* mice spent more time in nose contact and made more numerous and longer nose contacts with a cage mate versus the stranger mouse discovered on previous phase. Intranasal OT reversed this preference, resulting in longer time spent in nose contact (*G x D x S: F_1,47_=127.00, p<0.0001, p<0.0001)*, more in number (*G x D x S: F_1,47_=77.63, p<0.0001*) and length (*G x D x S: F_1,47_=136.62, p<0.0001*) contacts with the stranger mouse versus the cage mate. Vehicle-treated *Oprm1^-/-^* mice failed to display a preference during this phase; OT administration in these mutants led them to spend more time in nose contact and make longer contacts with the cage mate versus the stranger mouse. Thus, OT changed cage mate to stranger mouse preference in WT mice, while restoring preference for the cage mate in *Oprm1* null mice, as evidenced by social novelty preference ratio (*H_3,51_=37.15, p<0.0001*). Results are shown as scatter plots and mean ± sem. Solid stars: significant difference with the vehicle-treated *Oprm1^+/+^* group, Tuckey’s post-hoc test following a two-way ANOVA or 2-tailed t-test following a Kruskal-Wallis analysis of variance; open stars: genotype x treatment x stimulus interaction (stimulus: mouse/toy or stranger/cage mate comparison), Tukey’s post-hoc test following a repeated measure analysis of variance (ANOVA); daggers: genotype effect; one symbol: p<0.05, two symbols: p<0.01; three symbols: p<0.001. Letters: significant difference with vehicle-treated *Oprm1^-/-^* group (2-tailed t-test or Tukey’s post-hoc test); (c): p<0.05, (b): p<0.01, (a): p<0.001. More behavioural parameters in Fig. S2. C: cage mate, IU: International Units, M: mouse, OT: oxytocin, S: stranger, SI: social interaction, T: toy.

**Figure 2.**
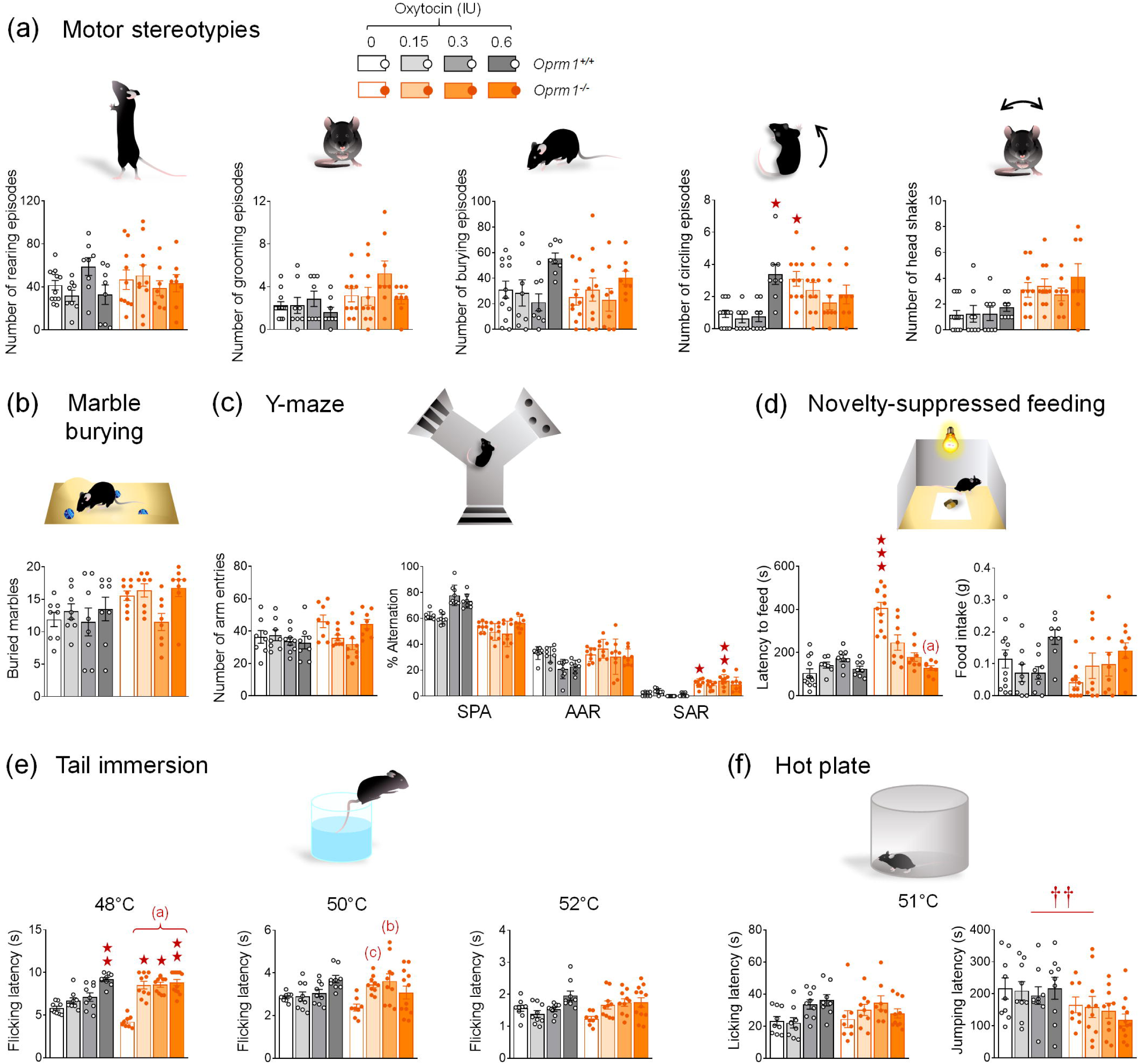
Acute per nasal OT relieved anxiety and induced analgesic effects in *Oprm1* null mice but had limited effects on stereotypies and perseveration. (a) When administered acutely 5 min before monitoring spontaneous motor stereotypies (*Oprm1^+/+^* vehicle: 6 males – 6 females, *Oprm1^-/-^*vehicle: 5 males - 5 females, *Oprm1^-/-^* OT 0.15 IU: 4 males – 6 females, other groups: 4 males – 4 females per genotype and dose), per nasal OT increased the number of circling events in *Oprm1^+/+^* mice. Vehicle-treated *Oprm1* null mice displayed more frequent circling behaviour, reduced (*H_7,72_=29.78, p<0.0001*) under OT administration. No effect of OT was detected on rearing, grooming or burying behaviour in mutant and wild-type mice, suggesting no effect on general activity. (b) No significant effect of genotype nor OT administration was detected in the marble burying test (*Oprm1^+/+^* OT 0.6 IU: 5 males – 3 females, *Oprm1^-/-^* vehicle: 5 males - 4 females, other groups: 4 males – 4 females per genotype and dose). (c) When exploring the Y-maze (*Oprm1^-/-^* OT 0.3 and 0.6 IU: 4 males – 5 females, other groups: 4 males – 4 females per genotype and dose), vehicle-treated and 0.3 IU OT-treated *Oprm1^-/-^* mice exhibited more frequent same arm returns than *Oprm1^+/+^* control mice *(H_7,66_ =24.1, p<0.01*); such perseverative behaviour did not reach significance in 0.15 and 0.6 OT-treated mutant mice. (d) In the novelty-suppressed-feeding test, latency to feed was increased in *Oprm1^-/-^* mice, while it did not differ from wild-type saline levels in OT-treated mice (*H_7,72_=39,8, p<0.0001*). (e) In the tail immersion test (*Oprm1^+/+^* groups: 4 males – 5 females, *Oprm1^-/-^* vehicle: 4 males – 4 females; *Oprm1^-/-^*OT at 0.15 and 0.5 IU: 5 males – 5 females, *Oprm1^-/-^* OT at 0.6 IU: 5 males – 7 females), OT-treated mice (0.6 IU in wild-type mice, all doses in *Oprm1* null mice) showed analgesia compared to saline-treated *Oprm1^+/+^* mice at 48°C (*H_7,76_=49.4, p<0.0001*). At 50°C, 0.15 and 0.3 IU of OT produced analgesic effects compared to saline treatment in *Oprm1^-/-^* mice (*H_7,76_=21.4, p<0.01*). At 52°C, OT increased nociceptive thresholds only in *Oprm1^-/-^*mice, at doses of 0.15 and 0.3 IU. No significant effect of OT was detected at 52°C. (f) In the hot plate test (same mice as for tail immersion), neither genotype nor OT treatment had a significant influence on nociceptive thresholds. Results are shown as scatter plots and mean ± sem. Solid stars: significant difference with the vehicle-treated *Oprm1^+/+^* group, Tuckey’s post-hoc test following a two-way ANOVA or 2-tailed t-test following a Kruskal-Wallis analysis of variance; daggers: genotype effect; one symbol: p<0.05, two symbols: p<0.01; three symbols: p<0.001. Letters: significant difference with vehicle-treated *Oprm1^-/-^* group (2-tailed t-test); (c): p<0.05, (a): p<0.001. AAR: alternate arm returns, MB: marble burying, SAR: same arm returns, SPA: spontaneous alternation.

**Figure 3.**
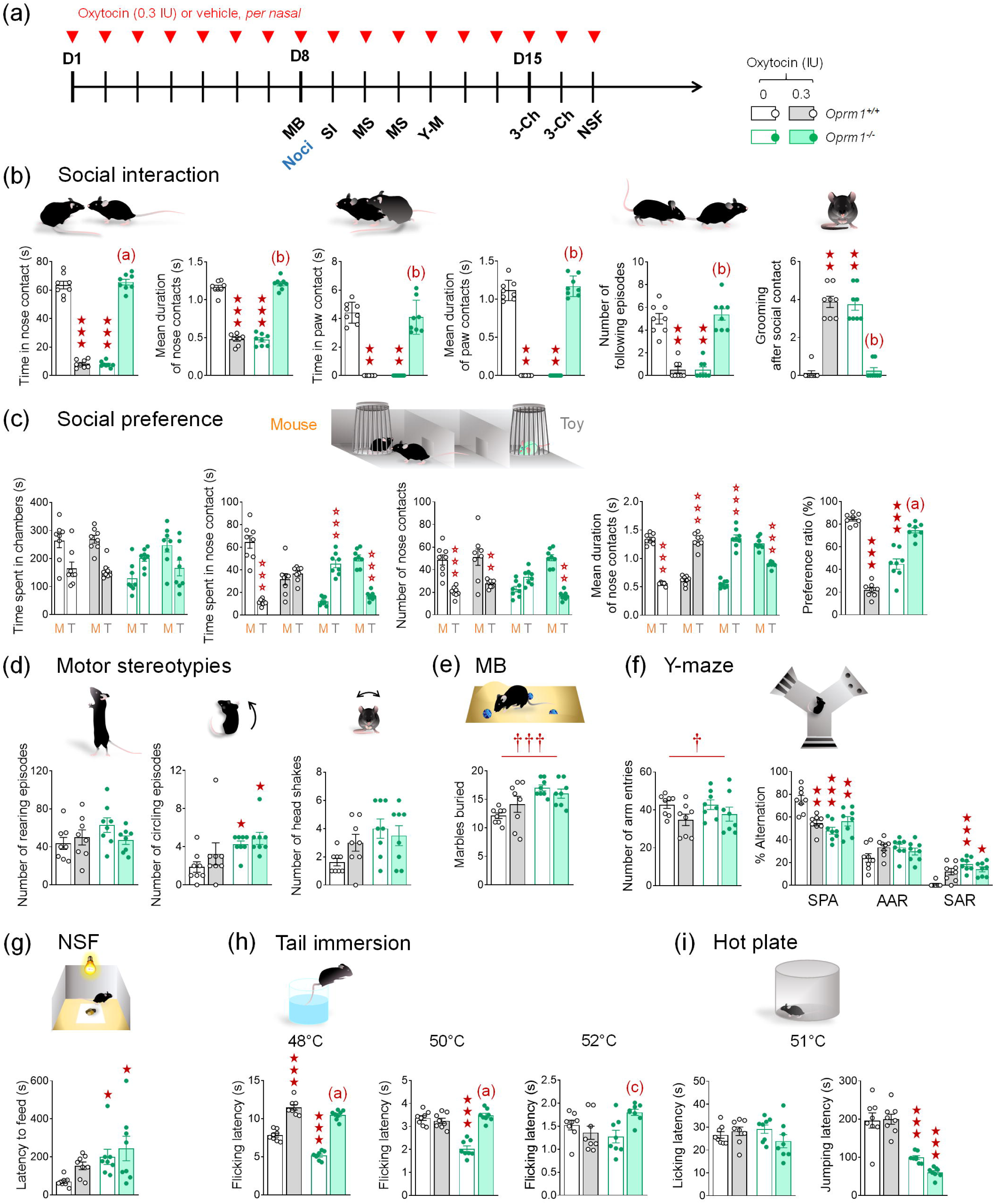
Chronic intranasal OT maintained beneficial on social behaviour in *Oprm1* knockout mice while had deleterious effects in wild-type controls. (a) A first cohort of *Oprm1^+/+^* and *Oprm1^-/-^* mice were treated daily with either OT (0.3 IU) or vehicle (4 males – 4 females per genotype and treatment) via per nasal route for 17 days. Behavioural testing started on D8. A second cohort received OT (0.3 IU) or vehicle (4 males – 4 females per genotype and treatment) daily for 8 days and was tested for nociception on D8. (b) In the direct social interaction test, social interaction was severely impaired in *Oprm1^+/+^*mice treated repeatedly with intranasal OT; conversely, OT restored the time spent in (*G x T: F_1,28_=994.1, p<0.0001*) and mean duration of (*G x T: F_1,28_=666.2, p<0.0001*) nose contacts, the time spent in (*H_3,32_=27.0, p<0.0001*) and mean duration of (*H_3,32_=26.8, p<0.0001*) paw contacts as well as the number of following episodes (*H_3,32_=24.3, p<0.0001*), and suppressing grooming after social contact (*H_3,32_=25.5, p<0.0001*) in *Oprm1^-/-^* mice. (c) Similarly, in the social preference test, repeated OT exposure compromised preference for the mouse over the toy in *Oprm1^+/+^* mice, but rescued preference for spending more time in (*G x T x S: F_1,28_=165.7, p<0.0001*), making more numerous (*G x T x S: F_1,28_=13.5, p<0.01*) and longer (*G x T x S: F_1,28_=789.8, p<0.0001*) nose contacts with the mouse versus the toy in *Oprm1* null mice. Resulting preference ratios were markedly decreased in the former while rescued in OT-treated *Oprm1^-/-^* mice (*G x T: F_1,28_=252.1, p<0.0001).* (d) *Oprm1^-/-^* mice display more frequent circling behaviour, and OT administration had no influence on this stereotyped behaviour (*H_3,32_=13.2, p<0.01*); no effect was detected in *Oprm1^+/+^* controls. (e) In the marble burying test, increased marble burying was observed in OT-treated as well as in vehicle-treated *Oprm1* knockouts (*H_3,32_=12.9, p<0.01*). (f) In the Y-maze, chronic OT failed to suppress perseverative same arm entries (SAR) in *Oprm1* mutants, and impaired spontaneous alternation (SPA) in *Oprm1^+/+^*mice (*G x T: F_1,28_=17.8, p<0.001).* (g) In the novelty-suppressed feeding test, *Oprm1* mutants took longer to eat in the middle of the arena, and OT treatment had no effect on this parameter (*H_3,32_=13.1, p<0.01*). (h) In the tail immersion test, nociceptive thresholds were significantly lower in *Oprm1^-/-^* mice and OT normalised them at 48°C while inducing analgesia in *Oprm1^+/+^*controls (*G x T: F_1,28_=10.3, p<0.01*). At 50°C, chronic OT normalised nociceptive thresholds in *Oprm1^-/-^* mice, without effects in WT mice (*G x T: F_1,28_=50.2, p<0.0001*). At 52°C, chronic OT increased the nociceptive threshold in *Oprm1* mutants (*H_3,32_=8.2, p<0.05*). (i) In the hot plate test, chronic OT failed to normalize lowered jumping latency in *Oprm1^-/-^* mice (*G: F_1,28_=91.2, p<0.0001*). Results are shown as scatter plots and mean ± sem. Solid stars: significant difference with the vehicle-treated *Oprm1^+/+^* group, Tuckey’s post-hoc test following a two-way ANOVA or 2-tailed t-test following a Kruskal-Wallis analysis of variance; open stars: genotype x treatment (Y-maze) or genotype x treatment x stimulus interaction (Social preference - stimulus: mouse/toy or stranger/cage mate comparison), Tukey’s post-hoc test following an analysis of variance (ANOVA); daggers: genotype effect; one symbol: p<0.05, two symbols: p<0.01; three symbols: p<0.001. Letters: significant difference with vehicle-treated *Oprm1^-/-^*group (2-tailed t-test or Tukey’s post-hoc test); (c): p<0.05, (b): p<0.01, (a): p<0.001. More behavioural parameters in Fig. S3. 3-Ch: 3-chamber social novelty test, AAR: alternate arm returns, D: day, MB: marble burying, MS: motor stereotypies, Noci: nociception, NSF: novelty-suppressed feeding, SAR: same arm returns, SPA: spontaneous alternation, Y-M: Y-maze.

#### Social abilities

##### Direct social interaction test

The experimental protocol was adapted from [57, 58]. On testing day, a pair of unfamiliar mice (not cage mates, age-, sex-, genotype- and treatment-matched) was introduced in one of 4 square arenas (50 x 50 cm, separated by 35 cm-high opaque grey Plexiglas walls) over a white infrared floor (View Point, Lyon, France) for 10 min (15 lx). Each arena received a black plastic floor (transparent to infrared) to minimize anxiety levels. The total amount of time spent in nose contact (nose-to-nose, nose-to-flank and nose-to-anogenital region), the number of these contacts, the time spent in paw contact and the number of these contacts, grooming episodes (allogrooming), notably ones occurring immediately (<5 s) after a social contact, as well as the number of following episodes were scored a posteriori on video recordings (infrared light-sensitive video camera) using an ethological keyboard (Labwatcher®, View Point, Lyon, France) by trained experimenters, and individually for each animal. The mean duration of nose and paw contacts was calculated as the number of events divided by the total time spent in these events.

##### Three-chamber social preference test

The experimental protocol was adapted from [57, 59]. The test apparatus consisted of a grey external Plexiglas box with transparent partitions dividing the box into three equal chambers (40 x 20 x 22.5 cm). Two sliding doors (8 x 5 cm) allowed transitions between chambers. Cylindrical wire cages (18 x 9 cm, 0.5 cm diameter-rods spaced 1 cm apart) were used to contain the mouse interactors and object (soft-toy mouse) placed in the two outward chambers of the 3-chamber social preference test. The test was performed under low-light conditions (15 lx) to reduce anxiety. Stimulus wild-type mice were habituated to wire cages for 2 days before the test (20 min/day). On testing day, the experimental mouse was introduced to the middle chamber and allowed to explore the whole apparatus for a 10-min habituation phase (wire cages empty). For the social preference phase, the experimental mouse was confined back in the middle-chamber while the experimenter introduced an unfamiliar wild-type age and sex-matched mouse (8-14-week-old, grouped housed) into a wire cage in one of the side-chambers and a soft toy mouse (8 x 10 cm) in the second wire cage. Then the experimental mouse was allowed to explore the apparatus for 10 min. For the modified novelty preference phase, the experimental mouse was returned to the middle chamber and the soft toy was replaced by a cage mate, to offer the choice to the experimental mouse of interacting whether with a very familiar mouse, the cage mate, or with the congener met during the first phase of the test. The sliding doors were reopened allowing an additional 10-min exploration. The time spent in each chamber and in nose contact with each wire cage, the number of these contacts and the number of entries in each chamber were scored a posteriori on video recordings using an ethological keyboard (Labwatcher®, View Point, Lyon, France) by trained experimenters. The mean duration of nose contacts (nose to nose, nose to flank, nose to anogenital region) was calculated from these data [57, 58]. Preference ratio was calculated as follows: Time in nose contact with the mouse / (Time in nose contact with the mouse + Time in nose contact with the object) x 100. The relative position of stimulus mice was counterbalanced between groups.

#### Stereotyped behaviours

##### Motor stereotypies

The experimental protocol was adapted from [60]. To detect spontaneous motor stereotypies in mutant versus wild-type animals, mice were individually placed in clear standard home cages (21×11×17 cm) filled with 3-cm deep fresh sawdust for 10 min. No water was available. Light intensity was set at 30 lux. Trained experimenters scored numbers of spontaneous head shakes, rearing, burying, self-grooming and circling episodes and the total amount of time spent burying by direct observation.

##### Marble-burying

Marble burying was used as a measure of stereotyped/perseverative behaviour [59, 61]. Mice were introduced individually in transparent cages (21×11×17 cm) containing 20 glass marbles (diameter: 1.5 cm) evenly spaced on 4-cm deep fresh sawdust. To prevent escapes, each cage was covered with a filtering lid. Light intensity in the room was set at 40 lx. The animals were removed from the cages after 15 min, and the number of marbles buried more than half in sawdust was recorded.

##### Y-maze exploration

Spontaneous alternation behaviour was used to assess perseverative behaviour [62, 63]. Each Y-maze (Imetronic, Pessac, France) consisted of three connected Plexiglas arms (15x15x17 cm) covered with distinct wall patterns (15 lx). Floors were covered with lightly sprayed fresh sawdust to limit anxiety. Each mouse was placed at the centre of a maze and allowed to freely explore this environment for 5 min. The pattern of entries into each arm was quoted on video-recordings. Spontaneous alternations (SPA), i.e. successive entries into each arm forming overlapping triplet sets, alternate arm returns (AAR) and same arm returns (SAR) were scored, and the percentage of SPA, AAR and SAR was calculated as following: total SPA or AAR or SAR / (total arm entries -2) * 100.

#### Anxiety-like behaviour

##### Novelty-suppressed feeding

The protocol was adapted from [64]. Novelty-suppressed feeding (NSF) was measured in 24-hr food-deprived mice, isolated in a standard housing cage for 30 min before individual testing. This test was performed in the same arenas as the ones used for direct social interaction. Three pellets of ordinary lab chow were placed on a white tissue in the centre of each arena, lit at 60 lx. Each mouse was placed in a corner of an arena and allowed to explore for a maximum of 15 min. Latency to feed was measured as the time necessary to bite a food pellet. Immediately after an eating event, the mouse was transferred back to home cage (free from cage-mates) and allowed to feed on lab chow for 5 min. Food consumption in the home cage was measured.

#### Nociceptive thresholds

##### Tail-immersion test

This test was performed as previously described [65, 66]. Nociceptive thresholds were assessed by immersing the tail of the mice (5 cm from the tip) successively into water baths at 48°C, 50°C and 52°C. Mice were gently maintained in a pocket during this experiment. The latency to withdraw the tail was measured at each temperature, with a cut-off of 10 s.

#### Oxytocin conditioning protocol

Animals were randomly distributed across four experimental conditions before behavioural assays had started: saline-object paradigm; OT 0.3 IU-object paradigm; saline-social interaction; OT 0.3 IU-social paradigm (see timeline in Figure 4).

**Figure 4.**
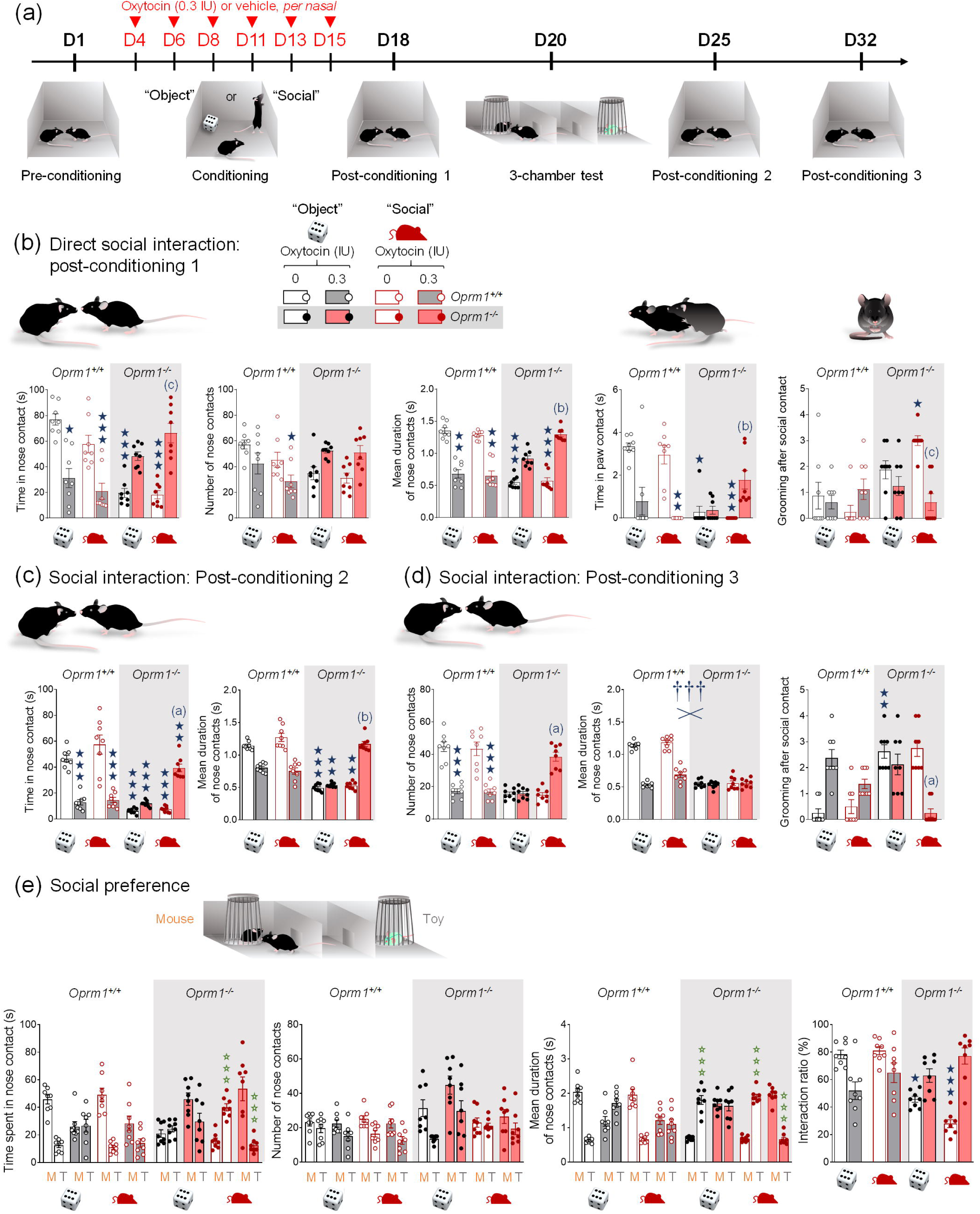
Beneficial effects of repeated intranasal OT on social deficit in *Oprm1* null mice were greater and lasted longer when associated with social experience. (a) After a pre-conditioning social interaction session, mice received per nasal OT (0.3 IU) or vehicle administration paired with the presentation of an unfamiliar object (“object” condition) or mouse (“social” condition) every two/three days over 2 weeks (D4 to D15) (4 males – 4 females per genotype, treatment and conditioning paradigm). A first post-conditioning social interaction session took place on D18, two days before 3-chamber test for social novelty preference (D20). Social interaction was assessed during two additional post-conditioning sessions, a week (D25) and two weeks (D32) after the first post-conditioning session. (b) During the first post-conditioning social interaction session, OT-treated *Oprm1^+/+^* mice displayed significant deficits in social behaviour, whereas OT-treated *Oprm1^-/-^* mice showed normalised social interaction parameters, as illustrated by opposite effects of OT treatment on the time spent in nose contact (*H_7,64_=41.4, p<0.0001*), duration of nose contacts (*H_7,64_=50.5, p<0.0001*) and time spent in paw contact (*H_7,64_=42.6, p<0.0001*). For these parameters, OT-treated *Oprm1^-/-^* mice tested under the social paradigm displayed significantly restored behaviour compared to vehicle-treated *Oprm1^-/-^*mice. The number of grooming events after social contact was normalised only in OT-treated *Oprm1^-/-^* mice tested under the social paradigm (*H_7,64_=27.3, p<0.001*). (c) After a week, detrimental effects of OT administration on the time spent in nose contact were still detectable in WT mice under the object paradigm, while beneficial effects in *Oprm1* knockouts were detected under the “social” paradigm only (*G x T X P: F_1,56_=15.6, p<0.001*). Regarding mean duration of nose contacts, deleterious effects of OT in *Oprm1^+/+^* mice did not reach significance; such duration was restored by OT in *Oprm1* null mice under the social condition only (*H_7,64_=54.6, p<0.0001*). (d) After another week, deleterious effects of OT in *Oprm1^+/+^* mice were detected on the number of nose contacts while beneficial effects of OT conditioning remained significant under the social paradigm only (*G x T X P: F_1,56_=10.9, p<0.01)*, as seen also for the number of grooming episodes after social contact (*H_7,64_=44.7, p<0.0001*). Regarding mean duration of nose contacts, OT decreased this parameter in *Oprm1^+/+^* mice while it had no detectable effect in *Oprm1* mutants (*G x T: F_1,56_=189,3, p<0.0001).* (e) In the three-chamber test, when the analysis included both *Oprm1^-/-^* mice and *Oprm1^+/+^* controls, we revealed that the time spent in nose contact with the mouse versus the object (*S x G x T: F_1,56_=74.6, p<0.0001)* and the mean duration of these contacts (*S x G x T: F_1,56_=207.0, p<0.0001)* were different depending on the genotype and OT exposure as well as depending on the genotype and conditioning paradigm (*S x T x P: F_1,56_=19.3, p<0.0001).* The number of nose contacts with each stimulus was under the influence of genotype and conditioning paradigm (*S x G x P: F_1,56_=5.8, p<0.05).* When focusing on mutant mice, we observed that preference for spending more time in nose contact (*S x T x P: F_1,28_=14.2, p<0.001)* and making longer nose contacts (*S x T x P: F_1,28_=27.8, p<0.0001)* with the mouse versus the toy were completely restored when *Oprm1^-/-^*mice were exposed to OT under the “social” but not the “object” paradigm. Finally, the social preference ratio was decreased in *Oprm1^-/-^* compared to *Oprm1^+/+^* mice and restored after OT exposure under both conditioning paradigms, although the contrast between vehicle and OT treatments was greater under the social paradigm (*H_7,64_=38.0, p<0.0001*). Results are shown as scatter plots and mean ± sem. Solid stars: significant difference with the vehicle-treated *Oprm1^+/+^*group, Tuckey’s post-hoc test following a two-way ANOVA or 2-tailed t-test following a Kruskal-Wallis analysis of variance; open stars: genotype x treatment (Y-maze) or genotype x treatment x stimulus interaction (Social preference - stimulus: mouse/toy or stranger/cage mate comparison), Tukey’s post-hoc test following an analysis of variance (ANOVA); daggers: genotype x treatment interaction; one symbol: p<0.05, two symbols: p<0.01; three symbols: p<0.001. Letters: significant difference with vehicle-treated *Oprm1^-/-^* group (2-tailed t-test or Tukey’s post-hoc test); (c): p<0.05, (b): p<0.01, (a): p<0.001. More behavioural parameters in Fig. S4. D: day, M: mouse, T: toy.

##### Pre-conditioning

Mice were evaluated for their basal social behaviour at day 1 (D1) of the protocol, using the direct social interaction test (10 min, protocol as described above).

##### Conditioning

Animals underwent 6 conditioning sessions at D4, 6, 8, 11, 13 and 15. During each conditioning session either OT (intranasal route, 0.3 IU) or saline were administered 5 min before a 10 min interacting session with either a novel mouse (age and sex-matched; “social” paradigm) or a novel object (“object” paradigm; dice, marble, Lego® brick, miniature plastic gem/rock/tree, wooden clip) each time, accordingly to the randomization group.

##### Post-conditioning

Drug-free mice undergone a direct social interaction test (10 min) at D18 and a three-chamber test for social preference (two phases of 10 min) at D20. To assess the maintenance of OT conditioning over time, two additional sessions of social interaction (10 min) were performed at D25 and 32.

A cohort of mice (social conditioning paradigm only) was dedicated to qRT-PCR analysis and sacrificed 45 min after the direct social interaction test on D18.

### 2.6 Real-time quantitative PCR analysis

Brains were removed and placed into a brain matrix (ASI Instruments, Warren, MI, USA). Caudate putamen (CPu), nucleus accumbens (NAc), ventral pallidum/olfactory tubercle (VP/Tu), lateral septum (LS) and central amygdala (CeA) were punched out while medial amygdala (MeA) was dissected from 1mm-thick slices (see Figure S1). Tissues were immediately frozen on dry ice and kept at -80°C until use. For each structure of interest, genotype and condition, samples were processed individually (n=8). RNA was extracted and purified using the Direct-Zol RNA MiniPrep kit (Zymo research, Irvine, USA). cDNA was synthesized using the ProtoScript II Reverse Transcriptase kit (New England BioLabs, Évry-Courcouronnes, France). qRT-PCR was performed in quadruplets on a CFX384 Touch Real-Time PCR Detection System (Biorad, Marnes-la-Coquette, France) using iQ-SYBR Green supermix (Bio-Rad) kit with 0.25 µl cDNA in a 12 µl final volume in Hard-Shell Thin-Wall 384-Well Skirted PCR Plates (Bio-rad). Gene-specific primers were designed using Primer3 software to obtain a 100- to 150-bp product and purchased from Sigma-Aldrich (Saint Quentin, France); sequences are displayed in Table S1. Relative expression ratios were normalised to the expression level of actin and the 2^−ΔΔCt^ method was applied to evaluate differential expression level. We focused primarily on genes coding for key players of the oxytocin/vasopressin system, peptides (*Oxt, Avp*) and receptors (*Oxtr, Avr1a, Avr1b*). We also measured the expression of marker genes of striatal projection neurons (SPNs) (*Drd1a, Pdyn, Tac1, Crh, Grm2, Drd2, Penk, Adora2, Grm4, Slc12a2, Slc12a5, Slc17a6, Slc17a7*) as well as markers of neuronal expression and plasticity (*Fos*, *Arc*), whose expression was found regulated in *Oprm1* knockout mice [45, 46, 49].

### 2.7 Statistics

Statistical analyses were performed using Statistica 9.0 software (StatSoft, Maisons-Alfort, France). For all comparisons, values of p<0.05 were considered as significant. Consistent with previous report [67], when conditions of normality were verified (Shapiro-Wilk test), statistical significance in behavioural experiments was assessed using one to four-way analysis of variance (treatment (T) or dose (D), genotype (G), and stimulus (S) effects) followed by Tukey’s multiple comparisons test. When these conditions were not fulfilled, we used the non-parametric Kruskal-Wallis analysis of variance followed by 2-tailed t-test to assess differences between groups. Under these conditions, genotype and treatment were collapsed in a single factor, and groups were analysed as independent. When a parameter was measured repeatedly (stimulus effect in the 3-chamber test: toy versus mouse, stranger versus cage-mate), however, non-parametric analysis would not allow post-hoc comparisons; thus, ANOVA was maintained, which may have exaggerated statistical significance. As male and female *Oprm1^-/-^* mice display similar behavioural deficits [45, 49] and preliminary experiments did not reveal differential OT effects between sexes, we pooled male and female data in the present study. As described previously [46, 67], qRT-PCR data were transformed prior to statistical analysis to obtain a symmetrical distribution centred on 0, using the following formula: if x<1, y=1-1/x; if x>1, y=x-1 (x: qPCR data; y: transformed data). Outliers over twice the standard deviation were excluded from calculations, as technical errors. Significance of qRT-PCR data was then assessed using a two-tailed t-test; an adjusted p value was calculated using Benjamini-Hochberg correction for multiple testing. Unsupervised clustering analysis was performed on transformed qRT-PCR data using complete linkage with correlation distance (Pearson correlation) for drug, treatment and brain region (Cluster 3.0 and Treeview software).

## 3 RESULTS

### 3.1 Acute *per nasal* administration of OT dose-dependently restored social behaviour in *Oprm1* null mice

We first assessed the effects of intranasal OT administration over a range of 3 doses: 0.15, 0.3 and 0.6 IU, on social behaviour in *Oprm1^-/-^* mice and their WT counterparts. When administered 5 min before behavioural testing, acute intranasal OT modified direct social interaction in *Oprm1* null mice following an inverted U-shaped dose response curve (Figure 1a). Indeed, vehicle-treated *Oprm1^-/-^* mice displayed a deficit in social interaction; this deficit was partially reversed after per nasal OT administration at 0.15 IU, fully at 0.3 IU but failed to be relieved at the dose of 0.6 IU. Deleterious effects of OT at 0.6 IU dose in *Oprm1^+/+^* mice were detected on the time spent in nose contact. Thus, the dose of 0.3 IU OT was the most efficient to restore social interaction in *Oprm1* null mice, when administered via per nasal route 5 min before testing. When administered 15 min before testing (Figure 1b), this dose of OT had partial effects on social interaction in mutant mice. When administered 30 min before testing (Figure 1c), acute intranasal 0.3 IU OT had no effect on social interaction. The delay of 5 min after nasal administration was thus used for the next experiments.

We then assessed whether OTR activation is indeed responsible for the effects of intranasal OT on social interaction in *Oprm1^-/-^* and *Oprm1^+/+^* mice. We injected the selective non-peptide OT antagonist LIT183 (pharmacological properties and chemical synthesis in Supplement 1) by intraperitoneal route at 7.5 or 15 mg/kg 25 min before intranasal OT (0.3 IU) administration and 30 min before testing (Figure 1d). In *Oprm1^-/-^* mice, OT failed to completely restore social interaction parameters when LIT183 was pre-administered. Thus, beneficial effects of intranasal OT on social interaction in *Oprm1^-/-^*mice relied on the activation of OT receptors, but likely not exclusively.

To further characterize the effects of acute intranasal OT in *Oprm1* null mice, we assessed the effects of this treatment on social preference and social novelty preference in the 3-chamber test. We used a modified version of the social novelty preference phase however, where the second stranger mouse was replaced by a cage mate. During the social preference phase (Figure 1e), intranasal OT fully rescued social preference in *Oprm1^-/-^* mice, with no effects in *Oprm1^+/+^* controls. During the modified social novelty preference phase (Figure 1f), WT mice displayed a preference for interacting with a cage mate versus the stranger mouse discovered on previous phase; intranasal OT completely reversed this preference. In contrast, *Oprm1* null mice failed to discriminate between cage mate and stranger mouse during this phase, and OT increased their interest for the cage mate. Thus, OT restored WT-like social preference and cage mate preference in *Oprm1^-/-^* mice.

### 3.2 Acute intranasal OT relieved anxiety and induced analgesic effects in *Oprm1* null mice but had little influence on stereotypies and perseveration

We then evaluated the effects of acute per nasal OT administration at 0.15, 0.3 or 0.6 IU on non-social behaviour in *Oprm1^-/-^*mice and their WT counterparts. Regarding spontaneous motor stereotypies (Figure 2a), *per nasal* OT reduced circling episodes in *Oprm1* knockout mice but increased this behaviour in *Oprm1^+/+^*mice (0.6 IU). No significant difference between groups was detected in the marble burying test (Figure 2b). When exploring the Y-maze (Figure 2c), saline and 0.3 IU OT-treated *Oprm1^-/-^* mice exhibited perseverative same arm returns (SAR). In the NSF test (Figure 2d), intranasal OT normalised normalised the latency to feed in *Oprm1^-/-^* mice.

In the tail immersion test (Figure 2e), per nasal OT showed analgesic properties at 0.15 and 0.3 IU in *Oprm1^-/-^* mice and at 0.6 IU in *Oprm1^+/+^* and *Oprm1^-/-^* mice at 48°C. At 50°C, per nasal OT showed analgesic properties at 0.15 and 0.3 IU in *Oprm1^-/-^*mice. No significant effect of OT was detected in the tail immersion test at 52°C nor in the hot plate test at 51°C (Figure 2f). Taken together, these results indicate that intranasal OT had little effect on stereotyped behaviour in *Oprm1* null mice but normalised their anxiety levels in the novelty-suppressed feeding test and induced analgesia in the tail immersion test.

### 3.3 Chronic intranasal OT remained beneficial on social behaviour in *Oprm1* knockout mice while exerted deleterious effects in wild-type controls

We then questioned whether beneficial effects of intranasal OT would maintain over repeated daily administration in *Oprm1^-/-^* mice. We administered OT via *per nasal* route at 0.3 IU once daily for 16 days and evaluated behaviour from day 8, starting behavioural tests 5 min after intranasal administration (Figure 3a).

Focusing first on social behaviour, we observed that repeated per nasal OT administration severely compromised social interaction in *Oprm1^+/+^* controls, leading to deficits of similar amplitude than those observed in vehicle-treated *Oprm1* mutants, but maintained its beneficial effects in *Oprm1^-/-^* mice (Figure 3b). Similarly, in the three-chamber test (Figure 3c), repeated OT administration dramatically impaired social preference in *Oprm1^+/+^* mice while maintaining its benefits in *Oprm1* mutants. Thus, chronic OT demonstrated prolonged beneficial effects on social behaviour in *Oprm1* null mice, whereas it was detrimental in WT mice.

Regarding motor stereotypies, chronic per nasal OT was ineffective reducing excessive spontaneous circling behaviour in *Oprm1^-/-^*mice (Figure 3d). Consistent with this, in the marble burying test, OT failed to normalise marble burying in *Oprm1* mutants (Figure 3e). In the Y-maze (Figure 3f), chronic OT failed to suppress perseverative same arm entries (SAR) in *Oprm1* mutants. In the novelty-suppressed feeding test (Figure 3g), chronic intranasal OT did not decrease feeding latency in *Oprm1^-/-^* mice. Together, these results indicate that repeated OT administration was not able to reduce stereotyped and perseverative behaviours or anxiety levels in *Oprm1^-/-^* mice.

We finally explored the analgesic effects of chronic intranasal OT in *Oprm1^-/-^* and *Oprm1^+/+^*mice. In the tail immersion test (Figure 3h), *Oprm1^-/-^*mice displayed lowered nociceptive thresholds that were normalised by OT at 48°C and 50°C. At 52°C, chronic OT increased the nociceptive threshold in *Oprm1* mutants. In the hot plate test (Figure 3i), chronic OT failed to normalize jumping latency in *Oprm1^-/-^* mice. Therefore, repeated OT administration maintained analgesic effects in the tail immersion test for both *Oprm1^-/-^* and *Oprm1^+/+^* mice, but was ineffective in the hot plate test at 51°C.

### 3.4 Beneficial effects of repeated intranasal OT on social deficit in *Oprm1* null mice were greater and lasted longer when associated with social experience

We challenged the social salience hypothesis by evaluating the influence of repeatedly pairing social experience to intranasal OT injection (social paradigm), compared to pairing with presentation of an inert novel object (object paradigm, Figure 4a). *Oprm1^-/-^* mice and their *Oprm1^+/+^* controls were tested for direct social interaction before receiving 6 administrations of OT (0.3 IU) or vehicle (every 2-3 days from D4 to D15) 5 min before entering an arena with an object or an unfamiliar conspecific. Social interaction was retested post-conditioning (drug free) on D18, D25 and D32. Social preference was evaluated on D20.

During preconditioning session, *Oprm1^-/-^* mice displayed a severe social deficit compared to *Oprm1^+/+^* mice (Figure S4a). In contrast, after repeated exposure to intranasal OT (D18, Figures 4b and S4b), social interaction was severely compromised in *Oprm1^+/+^* mice whilst social deficit was relieved in *Oprm1* knockouts. In mutant mice, however, the beneficial effects of OT were of higher amplitude when mice experienced social encounter immediately after OT administration (social paradigm). One week after first postconditioning assessment of social interaction (D25, Figures 4c and S4c), deficient social interaction was still detected in OT-treated *Oprm1^+/+^* mice trained under the “object” and “social” paradigms, while OT treatment maintained beneficial effects in *Oprm1^-/-^* mice only when trained under the “social” paradigm. After another week (D32, Figures 4d and S4d), detrimental effects of OT treatment were still detected for OT-treated *Oprm1^+/+^* mice trained under the “object” paradigm; beneficial effects of OT in *Oprm1^-/-^* mice trained under the “social” condition were not maintained for all parameters. Thus, beneficial effects of intranasal OT in *Oprm1* null mice were greater and longer lasting when this treatment was paired with social experience. The effects of repeated OT exposure were also assessed in the three-chamber test for social preference (D20, Figure 4e and S4e). To challenge the hypothesis that the conditioning paradigm influences the relieving effects of OT exposure on social preference in the *Oprm1* mouse model of ASD, we focused our analysis on *Oprm1^-/-^* mice and evidenced more significant restoration of social preference parameters under the “social” than the “object” paradigm.

In conclusion, repeated OT exposure better rescued social preference in *Oprm1* knockout mice when this treatment was associated with social experience.

### 3.5 Transcriptional consequences of social OT conditioning in *Oprm1* null mice and their wild-type controls

To gain insight into the molecular mechanisms at work in the brain of mice that underwent social OT conditioning, we assessed the effects of repeated OT exposure paired with social experience on gene expression 45 min after post-conditioning session (Figure 5a) in six regions of the reward/social circuit: CPu, NAc, VP/Tu, LS, MeA and CeA in *Oprm1^-/-^ and Oprm1^+/+^* mice. We focused on genes coding for key players of the oxytocin/vasopressin system, marker genes of SPNs and neuronal expression and plasticity. We monitored behaviour during post-conditioning session (Figure 5b) and confirmed previous observation of deleterious effects of intranasal OT exposure in *Oprm1^+/+^* mice contrasting with beneficial effects in *Oprm1^-/-^* mutants. We performed hierarchical clustering analysis of qRT-PCR data for each brain region to visualize the influence of OT conditioning on gene expression in *Oprm1^+/+^* and *Oprm1^-/-^* mice (Figure 5c). Transcriptional profiles were more similar between vehicle- and OT-treated *Oprm1^-/-^* mice in the NAc, VP/Tu, LS and CeA, showing predominance of genotype effects; OT treatment led to more similar profiles between OT-treated *Oprm1^+/+^* mice and OT-treated *Oprm1^-/-^*mice in the CPu and MeA. The main transcriptional effect of OT was to down-regulate gene expression across brain regions, as seen in the CPu (cluster1), NAc (cluster3), VP/TU (cluster2), MeA (cluster2) and CeA (cluster2), but not in the LS. Thus, OT treatment globally failed to normalize gene expression in *Oprm1* null mice.

**Figure 5.**
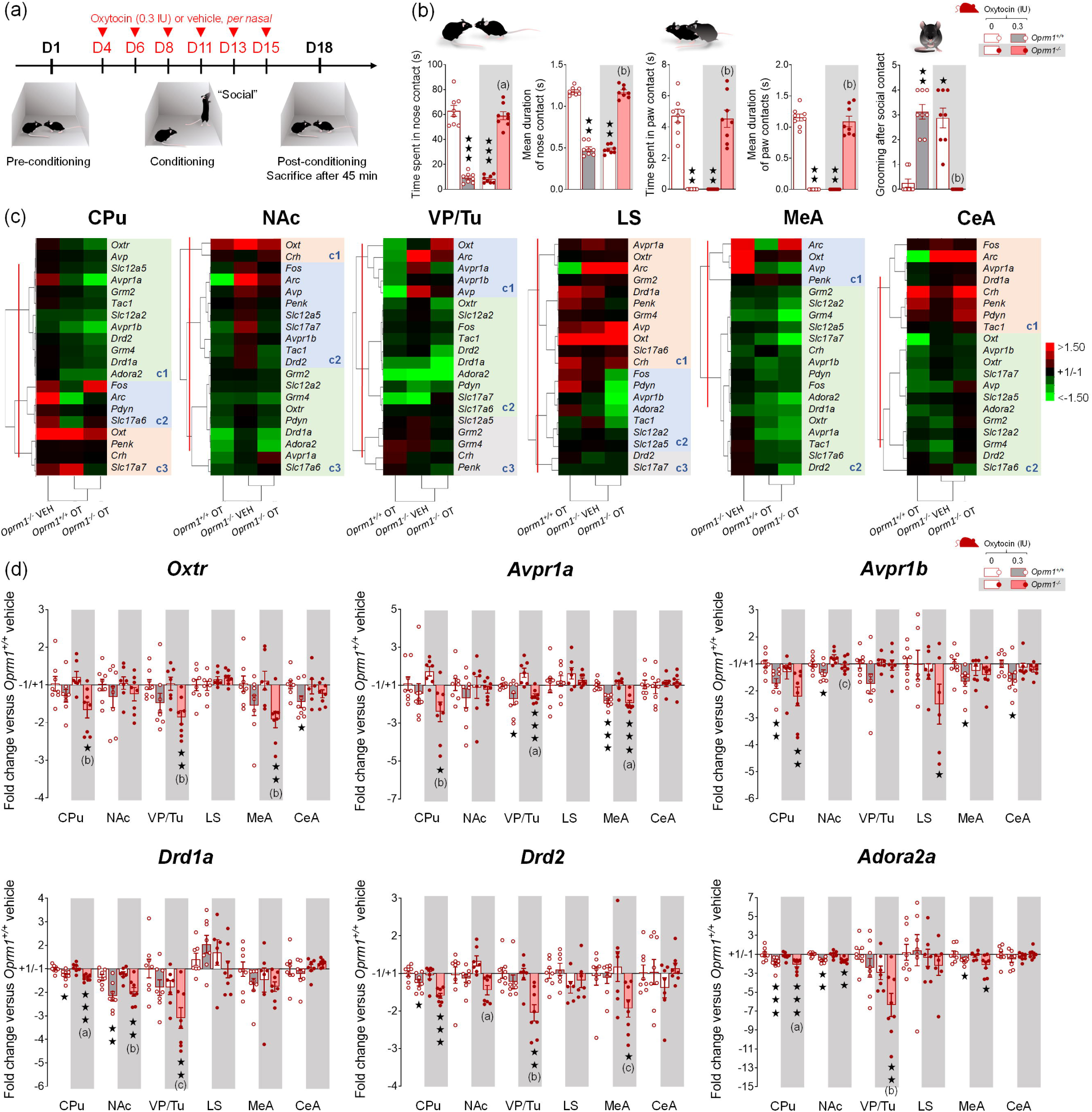
Transcriptional consequences of social OT conditioning in *Oprm1* null mice and their wild-type controls. (a) In this experiment, OT administration was paired with social encounter for all the mice (“social” paradigm; 4 males – 4 females per genotype and treatment). After a pre-conditioning social interaction session, mice received per nasal OT (0.3 IU) or vehicle administration paired with the presentation of an unfamiliar mouse every two/three days over 2 weeks (D4 to D15). The mice performed a post-conditioning social interaction session on D18 and were sacrificed 45 min after the beginning of behavioural assessment for qRT-PCR analysis. (b) As observed in previous experiment, OT exposure had opposite effects on social interaction in *Oprm1^+/+^*and *Oprm1^-/-^* mice, inducing a severe deficit in the former while rescuing interaction in the latter, as illustrated by the time spent in nose contact (*H_3,32_=23.3, p<0.0001*), mean duration of nose contacts (*H_3,32_=23.3, p<0.0001)*, time spent in paw contact (*H_3,32_=26.6, p<0.0001*) and duration of paw contacts (*H_3,32_=26.8, p<0.0001)*, as well as by the number of grooming episodes after a social contact (*H_3,32_=25.6, p<0.0001)* (more parameters in Fig. S5). (c) A hierarchical clustering analysis of qRT-PCR data was performed for each brain region of interest. The most contrasted transcriptional profiles were observed between OT-treated *Oprm1^+/+^* and OT-treated *Oprm1^-/-^*mice in the NAc, VP/Tu, LS and CeA, but not in the CPu and MeA where OT exposure led to more similar profiles between *Oprm1^+/+^* and *Oprm1^-/-^* mice. The main transcriptional effect of OT was to down-regulate gene expression across brain regions (green clusters), as seen in the CPu, NAc, VP/Tu, MeA and CeA, but not in the LS. (d) OT treatment decreased the expression of genes coding for oxytocin and vasopressin receptors (*Oxtr, Avpr1a, Avpr1b*) in the CPu, VP/Tu and MeA, more significantly in *Oprm1^-/-^*than in *Oprm1^+/+^* mice. Similarly, OT exposure led a down-regulation of the expression of the main marker genes of SPNs, the genes coding for the dopamine D1 (*Drd1a*) and D2 (*Drd2*) receptors, and the gene coding for the adenosine 2a (*Adora2*) receptor. Such down-regulation was more pronounced in the VP/Tu of the *Oprm1^-/-^*mice. Gene expression data are expressed as fold change versus *Oprm1^+/+^*- vehicle group (clustering or scatter plots and mean ± SEM). Comparison to *Oprm1^+/+^* - vehicle group (two-tailed t-test): One star p<0.05, two stars p<0.01, three stars p<0.001. Letters: significant difference with vehicle-treated *Oprm1^-/-^* group (2-tailed t-test); (c): p<0.05, (b): p<0.01, (a): p<0.001. qRT-PCR data used for clustering are displayed in Table S2. More individual transcriptional profiles for candidate genes are displayed in Fig. S5.

We then focused on candidate genes. We only took into consideration gene expression regulations affecting several brain regions for the same gene or several genes with similar functional profiles. Regarding the oxytocin/vasopressin system, transcriptome analysis revealed a global downregulation of the expression of genes coding for the OT (*Oxtr*) and vasopressin (*Avpr1a*, *Avpr1b*) receptors, in mice of both genotypes, after OT exposure (Figure 5d). This regulation affected mostly the CPu, VP/Tu and MeA, and was more significant in *Oprm1* knockouts than in WT controls. In contrast, OT treatment had little influence on the expression of genes coding for oxytocin and vasopressin (Table S2, Figure S5). Considering SPN markers, OT treatment decreased the expression of genes coding for the dopamine D1 (*Drd1a*) and D2 (*Drd2*) receptors, as well as the adenosine 2a (*Adora2*) receptor, in the CPu, NAc, and VP/Tu for the three genes, and also in the MeA for *Drd2* and *Adora2*. In the VP/Tu, down-regulation was more pronounced in *Oprm1* knockout mice compared to OT-treated WT mice. Thus, transcriptional results indicate that repeated OT exposure suppresses the expression of OT and vasopressin receptors, as well as the expression of the main SPN markers D1, D2 and A2a receptors, with a tendency for more pronounced effects in *Oprm1* null mice compared to wild-type mice.

## 4 DISCUSSION

The present study extends previous findings in *Oprm1* null mice [50, 51] by showing that beneficial effects of OT administration on social behaviour in *Oprm1^-/-^*mice tightly depend on the dose tested. Indeed, while administering OT acutely at doses commonly employed in animal and human studies, we observed an inverted U-shaped dose response curve on social interaction parameters, 0.15 IU being minimal, 0.3 IU producing optimal effects and 0.6 IU being deleterious. Detrimental effects of a high dose of OT may have resulted from excessive internalisation/uncoupling of OTR [51, 68], activation of vasopressin receptors [69] or recruitment of neural circuits involved in anxiety and fear [70-72]. In comparison, beneficial effects of OT have been detected from the dose of 200 µg/kg (0.075 IU) in mice prenatally exposed to sodium valproate or *Cntnap2* null mice [25, 29], suggesting that sensitivity to OT effects may vary between models.

Regarding kinetics, OT effects (0.3 IU) were optimal in *Oprm1^-/-^*mice at a short delay after administration (5 min), consistent with previous findings [40, 50], and vanished rapidly. Accordingly, OT is detected in the brain 5 min after intra-nasal delivery [19, 26]. After intraperitoneal administration, however, beneficial effects of OT on social behaviour in mouse models of ASD have been observed for up to 2 hours after administration [25, 29], likely due to persistent high blood levels of OT reached using this route [19]. In the present study, we focused on intranasal route as poorly invasive; it reveals, however, some limitations regarding duration of OT effects. As concerns frequency of administration, beneficial effects of OT on social interaction were maintained under repeated treatment in *Oprm1^-/-^*mice, as demonstrated in other ASD models [27, 30]. In contrast, such treatment severely and long-lastingly impaired social behaviour in wild-type mice, consistent with previous report [40]. Together, our results highlight the critical importance of the choice of dose and timing of OT administration for therapeutic use in the context of ASD. Moreover, a well-established ASD diagnostic appears a critical prerequisite to OT treatment, considering the detrimental behavioural effects of chronic OT in neurotypical subjects.

Improved social interaction following acute OT administration in *Oprm1^-/-^*mice involved OTR, as this effect was reduced in presence of the selective non-peptide OTR antagonist LIT183. However, a contribution of V_1A_ and V_1B_ vasopressin receptors is likely to account for the lack of complete reversal observed at a high dose of LIT183. Indeed, OT and vasopressin can bind to each other’s receptors to improve sociability [17, 73, 74].

We further explored OT effects in *Oprm1* null mice by assessing social preference. A single intranasal injection of OT restored preference for interacting with a congener over a mouse-shaped inert toy, likely by facilitating social approach [9, 75]. Interestingly, in *Oprm1* null mice, OT also restored preference for a familiar conspecific in our modified version of the social novelty phase, as observed in vehicle-treated *Oprm1*^+/+^ mice. Here, OT administration likely facilitated social memory and discrimination in *Oprm1* mutants [11, 32]. Conversely, OT administration reoriented social preference towards the most novel congener in *Oprm1*^+/+^ mice. This effect of OT in wild-type mice may be attributable to an attenuation of social fear or vigilance together with an increase in social approach, disinhibiting exploration of a novel conspecific [75-78]. Therefore, OT demonstrated beneficial effects on the social dimension of ASD-like deficits in *Oprm1* null mice, consistent with previous results in other murine models of ASD. Of note, the brain oxytocin system was found altered in all these models [26, 29, 79] as in *Oprm1* null mice, raising the hypothesis that oxytocin deficits are a prerequisite to successful OT treatment in ASD.

In our study, a single OT injection had little effect on the non-social dimension of ASD-like deficits in *Oprm1^-/-^* mice, namely stereotyped behaviours and perseveration, as previously reported for the *Cntnap2^-/-^* ASD mouse model [29]. However, improvements in stereotypies or cognitive flexibility have been observed in other models when OT administration was repeated [17, 80] or performed early in life [26, 79]. Similarly, in subjects with ASD, OT administration was either reported to decrease restricted/repetitive behaviour [81, 82] or to be inefficient on this dimension [33]. Such inconsistencies may reflect the recruitment of OT-sensitive brain substrates with antagonistic effects on repetitive behaviours, depending on the dose and frequency of administration [83, 84]. Consistent with anxiolytic properties [85], acute OT normalised anxiety levels in *Oprm1* mutants. This effect, however, was lost under repeated administration, maybe due to anxiogenic effects of OT developed under chronic administration [71]. In contrast, analgesic effects of OT on spinal nociception in *Oprm1^-/-^* mice were maintained upon chronic administration, suggesting the involvement of differential neuronal substrates [86, 87]. Thus, OT treatment appears more efficient on the core, social, dimension of ASD symptoms than stereotypies or secondary symptoms.

The main finding of our study was that the beneficial effects of OT on social behaviour in *Oprm1^-/-^* mice were greater and longer lasting when this neuropeptide was administered in a social context. These results are in line with a growing body of literature pointing towards social environment as key determinant for the prosocial effects of OT [42, 43]. OT therefore appears to behave as a coincidence detector for social experience and reward processes, allowing the reinforcement of social interactions. The neuronal substrate of such action would involve striatal regions, notably D_1_R and D_2_R-SPNs in the NAc, and interaction with multiple neuromodulators [9, 88]. In mice exposed to OT concomitantly with social experience, we found *Drd1a* and *Drd2* transcripts downregulated in the CPu, as well as in the NAc of *Oprm1^+/+^* and *Oprm1^-/-^* mice for the former, and in the VP/Tu of *Opmr1* null mice for the latter, pointing to modulation of dopaminergic transmission in these regions irrespective of genotype [89]. Interestingly, the expression of *Adora2a*, coding for adenosine A_2a_ receptors, was also consistently down-regulated in the striatum of *Oprm1* wild-type and mutant mice, and in the VP/Tu for *Opmr1* null mice. Such downregulation affected specifically D_2_R-expressing SPNs, of which *Adora2a* is a selective gene marker [90]. Not only A_2a_ receptors activate D_2_-SPNs but they also inhibit D_2_R signalling through heterodimerization [91]. Thus, decreased *Adora2a* expression may have facilitated a reduction in NAc D_2_R-SPN activity in *Oprm1* deficient mice, then facilitating social approach [67]. The contribution of D_1_R and D_2_R-SPNs to the social context-driven effects of OT deserves further exploration.

Interestingly, the expression of *Oxtr*, *Avpr1a* and *Avpr1b,* coding for OT and V_1a_ and V_1b_ vasopressin receptors, was also downregulated, a likely adaptive consequence of repeated exogenous OT administration. Decreased *Oxtr* expression in wild-type mice was previously reported after chronic intranasal OT (twice a day for 7 days) and proposed to contribute to the adverse effects of this treatment on social behaviour [40]. In our study, six OT injections were sufficient to trigger severe social deficit in *Oprm1^+/+^* mice but no marked regulation of *Oxtr* expression; in contrast, decreased *Oxtr* transcription was detected in *Oprm1^-/-^* mice, in which the benefits of chronic OT were preserved. Thus, *Oxtr* downregulation is unlikely to have contributed towards the deleterious or prosocial effect of OT in *Oprm1^+/+^*and *Oprm1^-/-^* mice, respectively. Deleterious effects OT in wild-type mice were more likely the consequence of long-term plastic rewiring in cortico-limbic circuits [92] and progressive recruitment of neural circuits involved in negative emotions [70-72].

In conclusion, our study provides several insights to better understand discrepancies in the results of recent clinical trials for OT in ASD [35]. First, this work highlights the crucial role of social context for OT effects. Consistent with our findings, when intranasal OT administration in children with ASD was immediately followed by positive social interaction, significant behavioural improvements were measured after a 6-week treatment, with the use of the gold standard ADOS-2 evaluation [93]. Together, these results strongly argue for combining OT administration with behavioural intervention [94]. Then, one may consider the use of a standard dose of OT in ASD as questionable and propose the re-evaluation of the therapeutic dose in future clinical studies. Different aetiologies, alterations in the OT system and/or reward circuit [95] as well as *OXTR* SNP variants may require adapting OT dose individually.

## ACKNOWLEDGMENT

We thank Audrey Léauté and Yannick Corde for their technical assistance. We thank the Experimental Unit PAO-1297 (EU0028, Animal Physiology Experimental Facility, DOI: 10.15454/1.5573896321728955E12) from the INRAE-Val de Loire Centre for animal breeding and care. We acknowledge the following funding sources: Région Centre (ARD2020 Biomédicament – GPCRAb), British Society for Neuroendocrinology (F Pantouli) and Marie-Curie/AgreenSkills Program (LP Pellissier). This work was supported by the Institut National de la Santé et de la Recherche Médicale (Inserm), Centre National de la Recherche Scientifique (CNRS), Institut National de Recherche pour l’Agriculture, l’Alimentation et l’Environnement (INRAe) and Université de Tours.

## DATA AVAILABILITY

Data will be made available on request.

## CRediT authorship contribution statement

**Fani Pantouli**: Formal analysis (equal); funding acquisition (supporting); methodology (supporting); visualization (supporting); writing—original draft (equal); writing—review and editing (equal); **Camille Pujol**: Formal analysis (equal); methodology (supporting); visualization (supporting); writing—original draft (equal); writing—review and editing (equal); **Cécile Derieux**: Data curation (supporting); formal analysis (equal); investigation (equal); methodology (equal); writing—review and editing (supporting); **Mathieu Fonteneau**: Data curation (supporting); formal analysis (equal); investigation (equal); methodology (supporting); writing—review and editing (equal); **Lucie P. Pellissier**: Conceptualization (supporting); funding acquisition (supporting); investigation (equal); methodology (supporting); writing— review and editing (supporting); **Claire Marsol**: Resources (supporting); methodology (supporting); writing—review and editing (supporting); **Julie Karpenko**: Resources (supporting); methodology (supporting); writing—review and editing (supporting); **Dominique Bonnet**: Resources (supporting); methodology (supporting); supervision (supporting); writing—review and editing (supporting); **Marcel Hibert**: Resources (supporting); methodology (supporting); supervision (supporting); writing—review and editing (supporting); **Alexis Bailey**: Conceptualization (supporting); funding acquisition (supporting); methodology (supporting); supervision (supporting); writing—review and editing (equal); **Julie Le Merrer**: Conceptualization (equal); data curation (equal); formal analysis (equal); funding acquisition (lead); investigation (supporting); methodology (equal); project administration (lead); visualization (lead); supervision (lead); writing—review and editing (equal); **Jerome AJ Becker**: Conceptualization (equal); data curation (equal); formal analysis (equal); funding acquisition (lead); investigation (supporting); methodology (equal); project administration (lead); visualization (supporting); supervision (lead); writing—review and editing (equal).

