## Supplementary for "Acute, chronic and conditioned effects of intranasal oxytocin in the mu opioid receptor knockout mouse model of autism: social context matters"

### LIT183, OTR antagonist

**Structure:**

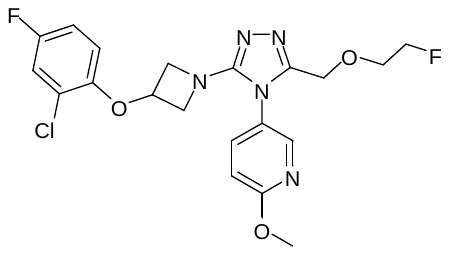

**Molecular mass:** 451.85 g/mol

**HR-MS (ES)** [M]: calculated 451.1223, observed 451.1235

**HPLC:** Tr = 4.33 min, HPLC purity: 100% at 220 nm. Gradient from 5 to 100% of CH_3_CN (0.1% TFA) in H_2_O (0.1 % TFA), 7.4 min at 1.6 mL/min, Ascentis Express C18 column (2.7 μm, 4.6 mm × 75 mm).

**logD (calc)** = 2.58

**Thermodynamic solubility:** 87.9 μg/mL in physiological serum (NaCl 0 .9%) at room temperature.

**Affinity** (TR-FRET, on HEK293 cells, n=3)^1^:

- OTR: Ki = 1.6 nM
- V1aR: Ki = 1364 nM
- V2R: Ki = 1469 nM

**Functional efficacy:**

|  | **Functional evaluation** | | | | **Notes** |
| --- | --- | --- | --- | --- | --- |
|  | **OXTR** | **V1aR** | **V1bR** | **V2R** |  |
| **PF3274167**  K_i_ | 9.5 | 1120 | >10000 | >10000 | Brown et al. 2010^2^ |
| **PF3274167**  IC_50_ | 8.9 ± 8.7 | 392 ± 41 | nd | nd | N=3, Calcium (Fluo4)^3^ |
| **LIT183**  IC_50_ | 3 ± 2 | 360 ± 340 | nd | nd | N=2, Calcium (Indo-1) ^3^ |
| **LIT183**  IC_50_ | 10 ± 6.6 | 390 ± 56 | nd | nd | N=3, Calcium (Fluo4) ^3^ |

nd: not detectable

**Synthesis:**

Synthetic scheme:

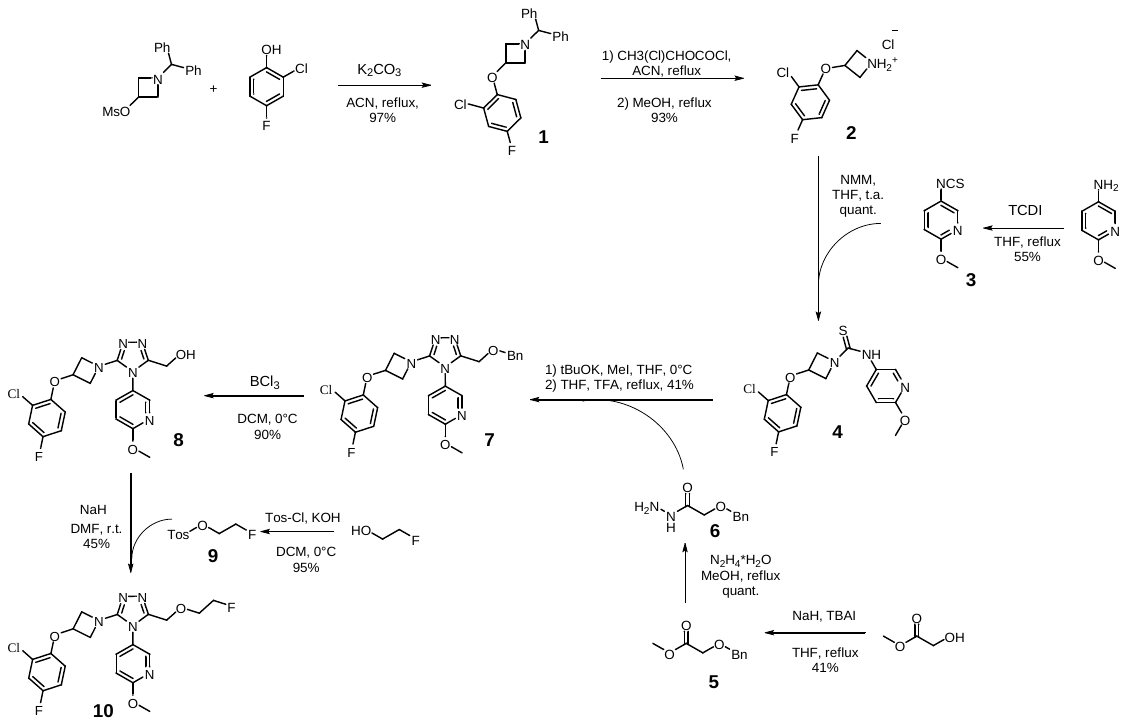

**Compound 1:^4^** To a solution of 1-(diphenylmethyl)azetidin-3-yl methanesulfonate (3.6 mmol, 1.14 g) and 2-chloro-4-fluorophenol (3.1 mmol, 460 mg) in dry ACN (20 mL) K_2_CO_3_ (7.5 mmol, 1.08 g) is added. The reaction mixture is stirred under reflux for 4 h. Water (150 mL) is added and the resulting mixture is extracted with CH_2_Cl_2_ (150 mL). The organic layer is washed with water (2x150 mL), brine (150 mL) and the volatiles are removed under vacuum. The residue is purified by reverse phase flash chromatography eluted with ACN/H_2_O (5% to 100% in 30 min) to give **1** as yellow oil (1.1g, 97%).

**Compound 2:^5^** To a stirred solution of **1** (0.87 mmol, 320 mg) in dry CH_3_CN (5 mL) at 0°C 1-chloroethyl chloroformate (1.74 mmol, 188 µL) is added dropwise. The reaction mixture is stirred at 80°C for 1h monitored by HPLC then cooled down to r.t. and concentrated under reduced pressure. The residue is dissolved in dry MeOH (5 mL) and stirred at 65°C for 3h monitored by HPLC. Water (50 mL) is added and the solution is washed with heptane (3x50 mL), then the volatiles are removed under vacuum to give **2** as white solid (193 mg, 93%).

**Compound 3:^6^** A solution of 5-amino-2-methoxypyridine (0.5 mmol, 40.2 µL) in dry THF (3 mL) is added dropwise over the period of 40 min to a stirred solution of 1,1'-thiocarbonyldiimidazole (0.75 mmol, 148.5 mg) in dry THF (2 mL) cooled to ice-water temperature. The reaction mixture is allowed to warm to room temperature then stirred for 30 min. THF is evaporated, the residue is dissolved in CH_2_Cl_2_, washed with NaHCO_3_ (sat), brine then concentrated under vacuum and filtered through a silica gel pad, washed with CH_2_Cl_2_. The solvent is removed under vacuum to give **3** as white solid (46 mg, 55%).

**Compound 4:** To a suspension of **2** (3 mmol, 707 mg) in dry THF (15 mL) at 0°C, NMM (3.6 mmol, 392 µL) is added. A solution of **3** (3 mmol, 494 mg) in dry THF (15 mL) is added dropwise. The reaction mixture is heated to r.t. and stirred for 1h. The resulted mixture is concentrated under vacuum. To the residue DCM (150 mL) is added and the organic fraction is washed with water (3x150 mL), brine (150 mL), dried over Na_2_SO_4_. The volatiles are removed under vacuum to give **4** as white powder (1.1 g, quantitative).

**Compound 53:^4^** To a solution of methyl glycolate (5 mmol, 396 µL) in dry THF (25 mL) under Ar at 0°C, NaH (5.25 mmol, 60% dispersion in oil, 210 mg) is added. The resulted mixture is stirred at 0°C for 30 min, then NBu_4_I (0.5 mmol, 185 mg) is added followed by BnBr (5 mmol, 610 µL). The reaction mixture is heated at 60°C for 24h. Water (100 mL) is added and the mixture is extracted with CH_2_Cl_2_ (2x50 mL). The combined organic fractions are washed with water (2x100 mL) and brine (100 mL). The volatiles are removed under vacuum and the residue is purified consequently by flash chromatography eluted with EA/heptane (5% for 5 min, then from 5% to 40% in 15 min) and by reverse phase flash chromatography eluted with ACN/H_2_O (from 20% to 100% over 30 min) to give **5** as yellow oil (41%).

**Compound 6:** To a solution of **5** (2.5 mmol, 450 mg) in dry MeOH (20 mL) hydrazine monohydrate (3.7 mmol, 185 µL) is added. The reaction mixture is refluxed for 6h, then the volatiles are removed under vacuum, co-evaporated 3 times with new portions of MeOH, the residue is dried under high vacuum to give the desired compound as colourless oil (448 mg, quantitative).

**Compound 7:** To a stirred solution of **4** (1.2 mmol, 441 mg) in dry THF (17 mL) cooled to ice-water temperature tBuOK (1.44 mmol, 170 mg) is added. The resulting mixture is stirred for 5 min then MeI (1.44 mmol, 90 µL) is added. The reaction mixture is stirred at ice-water temperature for 15 min. Water (150 mL) is added and the resulted mixture is extracted with EA (2x70 mL). The combined organic fractions are washed with water (3x150 mL), brine (150 mL), dried over Na_2_SO_4_ and the volatiles are removed under vacuum. The residue is dissolved in dry THF (17 mL). To the obtained solution **6** (1.43 mmol, 257 mg) is added followed by TFA (0.59 mmol, 44 µL). The reaction mixture is stirred under reflux for 6h followed by HPLC, then the volatiles are removed under vacuum. To the residue water (100 mL) is added and the mixture is extracted with DCM (2x50 mL). The organic layer is washed with water (2x100 mL), brine (100 mL) and dried over Na_2_SO_4_. The volatiles are removed under vacuum and the residue is purified by reverse phase flash chromatography eluted with ACN (with 0.1% TFA) / H_2_O (with 0.1% TFA) (5-100% over 30 min) to give **7** (300 mg, 41%) as yellow oil.

**Compound 8:** To a solution of **7** (0.11 mmol, 68 mg) in dry DCM (5 mL) cooled to ice-water temperature a solution of BCl_3_ in DCM (1 M, 0.56 mmol, 56 µL) is added. The resulting mixture is stirred for 30 min. Water (50 mL) and DCM (50 mL) are added. A solution of 10N NaOH was added dropwise until the pH of the aqueous phase was 13. The organic fraction is separated, washed with 1N solution of NaHCO_3_ (50 mL), brine (50 mL), dried over Na_2_SO_4_ and the volatiles are removed under vacuum. The residue is purified by reverse phase flash chromatography eluted with ACN/H_2_O (5% to 100 % in 30 min) to give after lyophilization compound **8** as yellow solid (230 mg, 90%).

**Compound 9:** To a solution of 2-fluoroethan-1-ol (0.5 mmol, 29 µL) in dry DCM (5 mL) cooled to ice-water temperature KOH (1.5 mmol, 84 mg) and Tos-Cl (0.55 mmol, 107 mg) are added. The reaction mixture is stirred at 0°C for 2h then at r.t. overnight. Water (100 mL) and DCM (50 mL) are added. The organic fraction is separated, washed with water (2x50 mL), brine (50 mL), dried over Na_2_SO_4_ and the volatiles are removed under vacuum. The residue is purified by flash chromatography eluted with EA/heptane (5% - 100% over 30 min) to give **9** as colourless oil (105 mg, 96%).

**Compound 10:** To a solution of **8** (0.038 mmol, 20 mg) in dry DMF (1 mL) under Ar, KOH (0.19 mmol, 11 mg) is added. The resulted mixture is stirred at 0°C for 10 min, then **9** (0.058 mmol, 12.6 mg) is added. The reaction mixture is stirred at ice-water temperature for 3h, then at r.t. overnight. Water (100 mL) and DCM (50 mL) are added. The organic fraction is separated, washed with water (2x50 mL), brine (50 mL), dried over Na_2_SO_4_ and the volatiles are removed under vacuum. The residue is purified by reverse phase flash chromatography eluted with ACN/H_2_O (5% to 100 % in 30 min) to give after the lyophilization **10** as white solid (15 mg, 69%).

### Supplementary figures

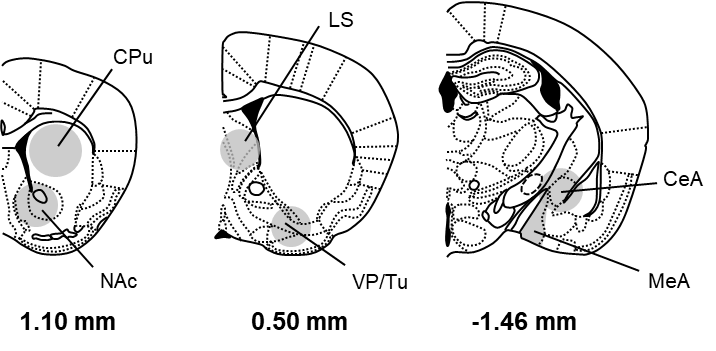

**Figure S1. Schematic representations depict brain regions dissected for gene expression study**. CPu, NAc, LS, VP/Tu, MeA and CeA, were punched on 1-mm thick brain slices (CPu: one punch/side, ᴓ 2 mm; NAc, BNST, VP, CeA, and VTA/SNc: one punch/side, ᴓ 1.25 mm). Coordinates refer to bregma. CPu: Caudate Putamen; CeA: Central Amygdala; LS: lateral septum; NAc: Nucleus Accumbens; VP/Tu: ventral pallidum/olfactory tubercle.

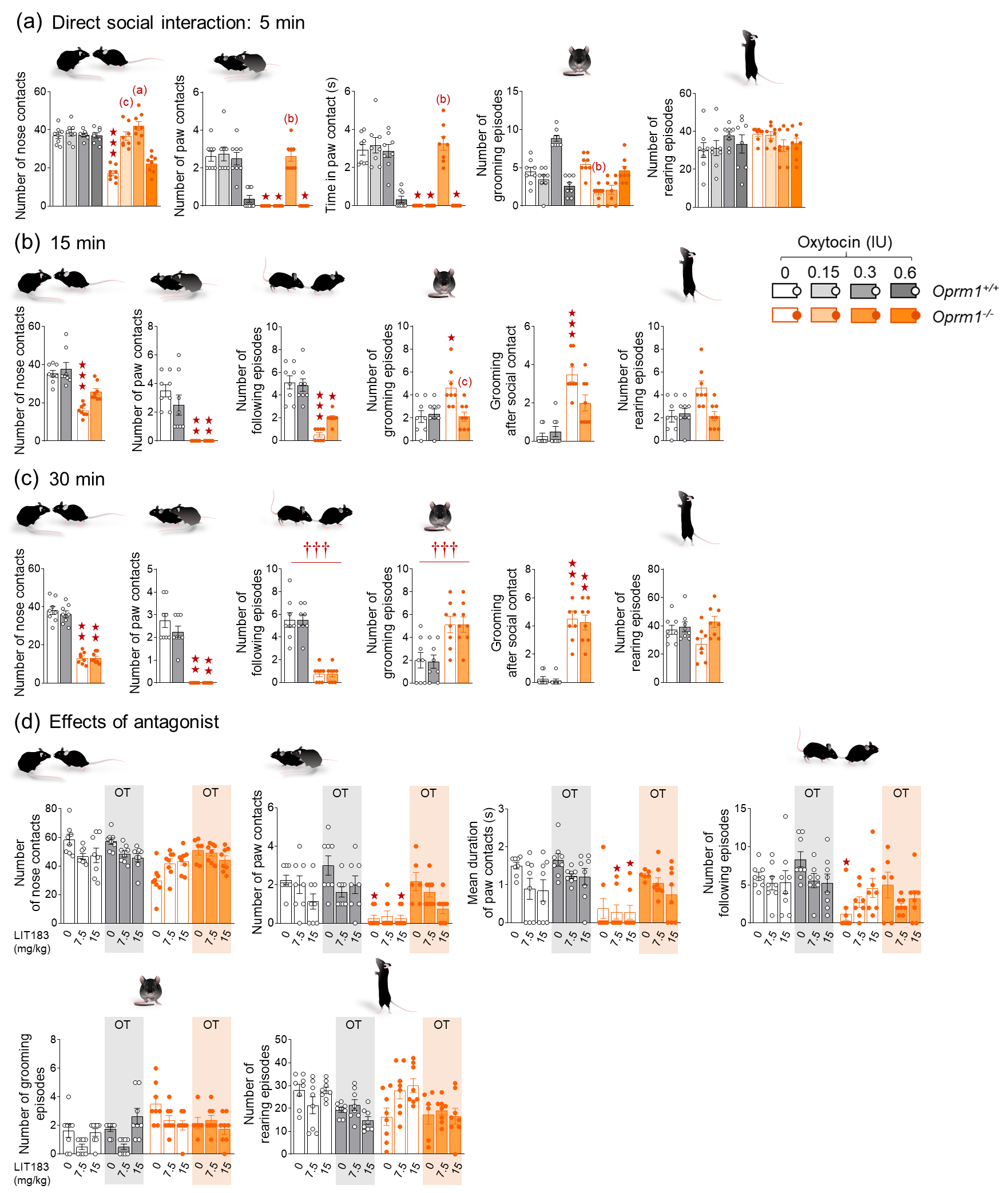

**Figure S2. Acute per nasal administration of OT dose-dependently restored social behaviour in *Oprm1* null mice.** **(a)** *Oprm1^+/+^* and *Oprm1^-/-^* mice received OT or vehicle (4 males – 4 females per genotype and treatment) via per nasal route 5 min before the direct social interaction test at the dose of 0, 0.15, 0.3 or 0.6 IU. Vehicle-treated *Oprm1^-/-^* mice displayed a severe deficit in social interaction parameters ; OT at 0.15 and 0.3 IU fully restored the number of nose contacts (*Genotype [G] x Dose [D] interaction:* *F_1,56_=22.19, p<0.0001*) while only the dose of 0.3 IU restored the number of paw contacts (*H_7,64_=53.72, p<0.0001*) and time spent in (*H_7,64_=52.71, p<0.0001*) paw contact in mutant mice. Moreover, OT at 0.3 IU decreased the number of grooming episodes in *Oprm1^-/-^* mice (*H_7,64_=42.15, p<0.0001).* **(b)** When administered 15 min before testing (4 males – 4 females per genotype and treatment), the optimal dose of 0.3 IU OT restored the number of nose contacts in *Oprm1* null mice (*H_3,32_=23.67, p<0.0001)* but failed to normalize their number of paw contacts (*H_3,32_=27.27, p<0.0001)* and following episodes (*H_3,32_=25.89, p<0.0001)*. However, OT reduced the number of grooming episodes (*H_3,32_=11.60, p<0.01)* and grooming after social contact in mutant mice (*H_3,32_=22.27, p=0.0001)*. **(c)** When administered 30 min before testing (4 males – 4 females per genotype and treatment), per nasal OT at 0.3 IU failed to relieve social interaction deficit in *Oprm1* null mice. **(d)** The non-peptide OT antagonist LIT183 or its vehicle (doses of 0, 7.5 or 15 mg/kg) were administered intraperitoneally 25 min before per nasal OT administration (0.3 IU) and 30 min before direct social interaction test (4 males – 4 females per genotype, LIT183 doses and OT treatment). *Oprm1* null mice treated with intranasal vehicle displayed reduced number (*H_11,94_=47.13, p<0.0001)* and duration of paw contacts (*H_11,94_=36.27, p<0.0001)* that were not detected in *Oprm1* null mice treated with intranasal OT, independently from LIT183 administration. *Oprm1^-/-^* mice of the vehicle/vehicle group displayed a reduced number of following episodes (*H_11,94_=32.92, p<0.001)*, not significantly detected in other treatment groups.

Results are shown as scatter plots and mean ± sem. Solid stars: significant difference with the vehicle-treated *Oprm1^+/+^* group, Tuckey’s post-hoc test following a two-way ANOVA or 2-tailed t-test following a Kruskal-Wallis analysis of variance; one symbol: p<0.05, two symbols: p<0.01; three symbols: p<0.001. Letters: significant difference with vehicle-treated *Oprm1^-/-^* group (2-tailed t-test or Tukey’s post-hoc test); (c): p<0.05, (b): p<0.01, (a): p<0.001. IU: International Units, OT: oxytocin.

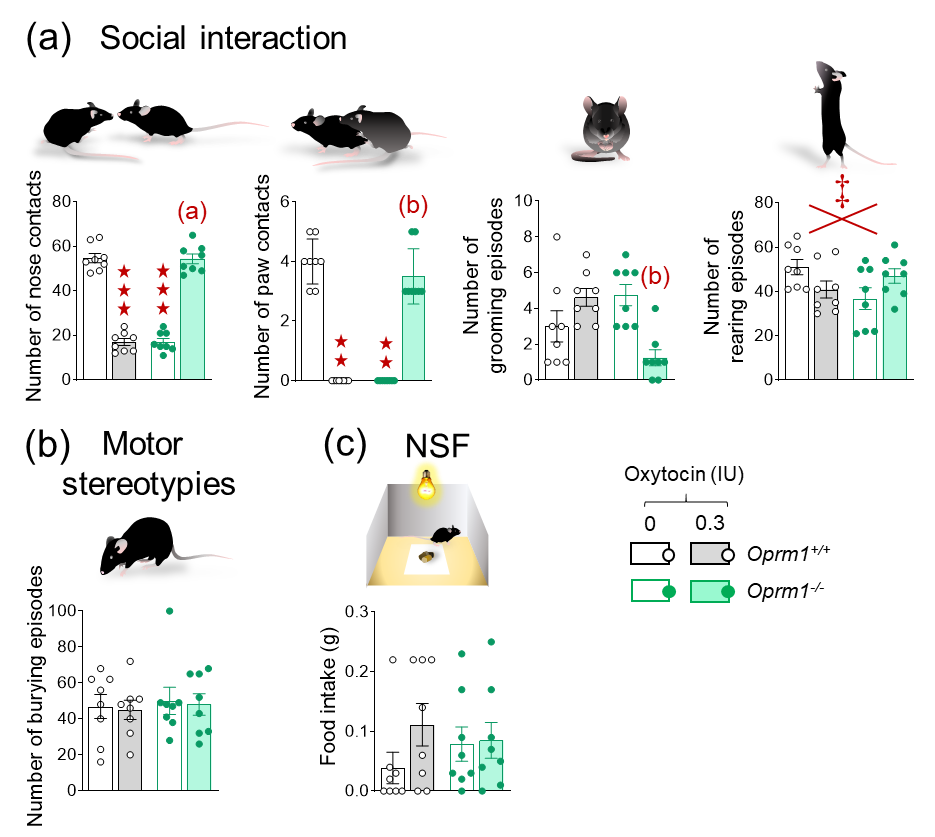

**Figure S3. Chronic per nasal administration of OT at 0.3IU restored social interaction and suppressed stereotypies and anxiety-like behaviour in *Oprm1* null mice but had deleterious effects in the same behaviours in their WT counterparts.** *Oprm1^+/+^* and *Oprm1^-/-^* mice received 0.3IU of OT or vehicle (4 males – 4 females per genotype and treatment) once a day for 17 consecutive days, 5 min before testing (timeline in Figure 3a). **(a)** Vehicle-treated *Oprm1^-/-^* mice displayed a severe deficit in social interaction parameters, such as number of nose (*F_1,28_=405.80; p<0.0001*) and paw (*H_3,32_=27.66, p<0.001*) contacts; OT treatment fully reversed these deficits. Chronic OT at 0.3 IU decreased the number of grooming episodes in *Oprm1^-/-^* mice (*H_3,32_=14.53, p<0.01)* and tended to oppositely modulate the number of rearing episodes (*F_1,28_=6.77; p<0.05*). Neither genotype nor OT treatment significantly modified **(b)** the number of burying episodes when assessing spontaneous motor stereotypies or **(c)** the amount of food consumed following the NSF test.

Results are shown as scatter plots and mean ± sem. Solid stars: significant difference with the vehicle-treated *Oprm1^+/+^* group, Tuckey’s post-hoc test following a two-way ANOVA or 2-tailed t-test following a Kruskal-Wallis analysis of variance; one symbol: p<0.05, two symbols: p<0.01; three symbols: p<0.001. Letters: significant difference with vehicle-treated *Oprm1^-/-^* group (2-tailed t-test or Tukey’s post-hoc test); (b): p<0.01, (a): p<0.001. IU: International Units, NSF: novelty-suppressed feeding test.

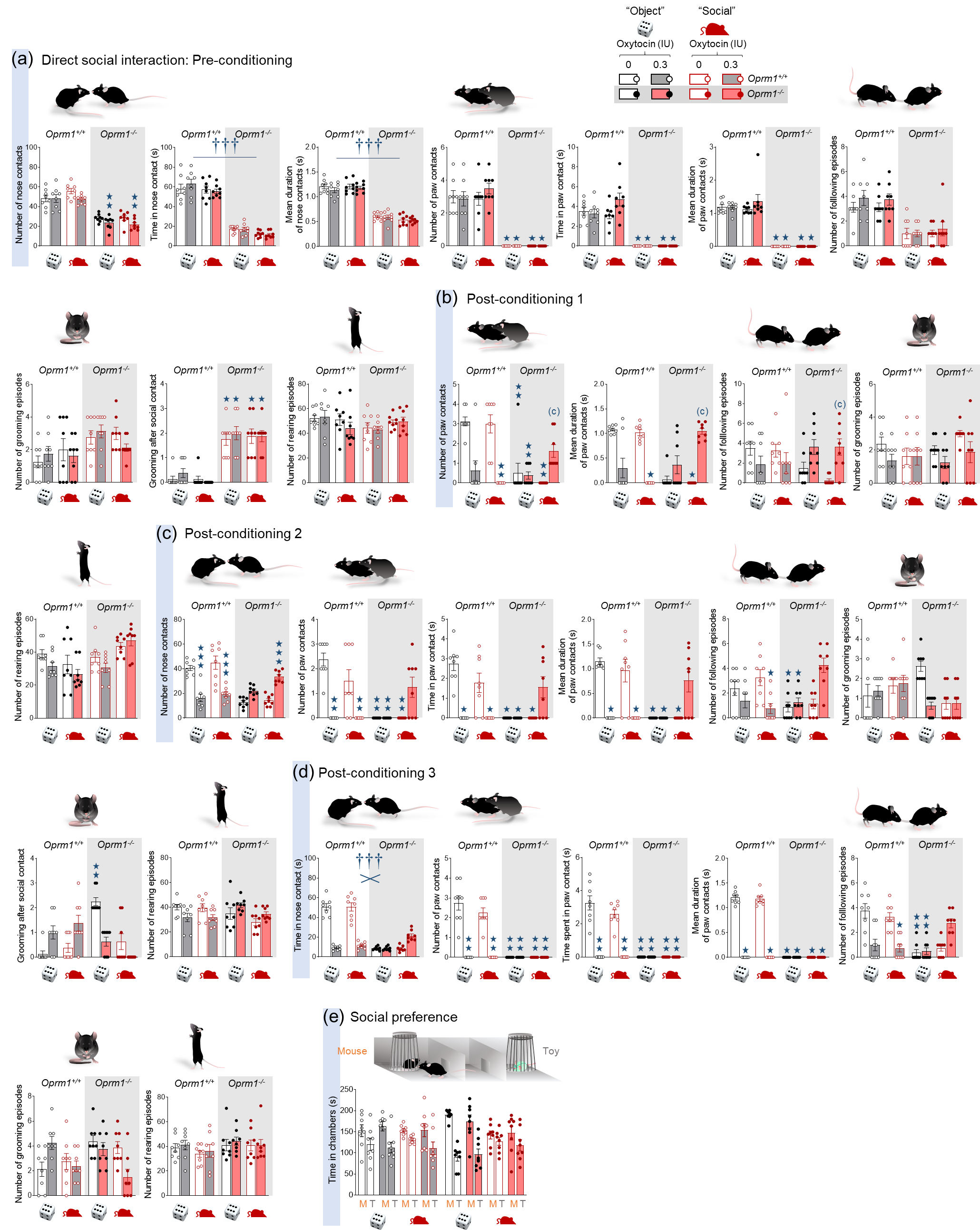

**Figure S4. Beneficial effects of repeated intranasal OT on social deficit in *Oprm1* null mice were greater and lasted longer when associated with social experience.** After a pre-conditioning social interaction session, mice received per nasal OT (0.3 IU) or vehicle administration paired with the presentation of an unfamiliar object (“object” condition) or mouse (“social” condition) every two/three days over 2 weeks (D4 to D15) (4 males – 4 females per genotype, treatment and conditioning paradigm). A first post-conditioning social interaction session took place on D18, two days before 3-chamber test for social novelty preference (D20). Social interaction was assessed during two additional post-conditioning sessions, a week (D25) and two weeks (D32) after the first post-conditioning session (timeline in Figure 4a). **(a)** During the pre-conditioning social interaction session (no OT exposure yet), *Oprm1^-/-^* mice showed significant deficits in social behaviour compared to *Oprm1^+/+^* mice, as illustrated by decreased number of nose (*G:* *F_1,56_=9481.89; p<0.0001*) and paw contacts (*H_7,64_=55.36, p<0.0001*), time spent in nose (*H_7,64_=50.38, p<0.0001*) and paw contacts (*H_7,64_=55.42, p<0.0001*) and their duration (nose: *G:* *F_1,56_=482.00; p<0.0001;* paw: *H_7,64_=55.8, p<0.0001*), as well as a decreased number of following episodes (*H_7,64_=37.04, p<0.0001*). In addition, *Oprm1^-/-^* mice showed an increased number of grooming episodes after a social contact, but no difference in the total number of grooming episodes nor the number of rearing episodes. **(b)** During the first post-conditioning social interaction session, OT-treated *Oprm1^+/+^* mice displayed significant decrease in the number (*H_7,64_=44.7, p<0.0001*) and mean duration of paw contacts (*H_7,64_=37.4, p<0.0001*) while in OT-treated *Oprm1^-/-^* mice these parameters were improved under the social paradigm only. Similarly, the number of following episodes was increased in OT-treated *Oprm1^-/-^* mice under the social setting (*H_7,64_=22.2, p<0.0001*). Neither genotype, treatment nor experimental paradigm had a significant influence on the number of grooming episodes in this test. OT-treated *Oprm1^-/-^* mice under the social setting tended to display more frequent rearing episodes (*H_7,64_=22.3, p<0.05*). **(c)** During the second post-conditioning social interaction session (D25), social interaction parameters remained severely impaired in OT-treated *Oprm1^+/+^* mice while significant effects of OT exposure could still be detected in OT-treated *Oprm1^-/-^* mice under the social paradigm, as observed for time spent in paw contact(*H_7,64_=45.9, p<0.0001*), the number of such contacts (*H_7,64_=46.5, p<0.0001*) and their mean duration (*H_7,64_=44.9, p<0.0001*), the number of following episodes (*H_7,64_=24.8, p<0.001*) and the number of grooming episodes after social contact (*H_7,64_=36.06, p<0.0001*). Neither genotype, treatment nor experimental paradigm had a significant influence on the number of grooming nor rearing episodes in this test. **(d)** After an additional week (D32), OT decreased the time spent in nose contact (*G x T:* *F_1,56_=242.28, p<0.0001)*, the number of paw contacts (*H_7,64_=61.6, p<0.0001*), time spent in paw contact (*H_7,64_=61.4, p<0.0001*) and mean duration of paw contacts (*H_7,64_=61.3, p<0.0001*) in *Oprm1^+/+^* mice while it had no detectable effect in *Oprm1* mutants. In contrast, the number of following episodes was significantly reduced only in OT-treated *Oprm1^+/+^* mice tested under the “social” setting, and in *Oprm1^-/-^* mice tested under the “object” setting (*H_7,64_=38.5, p<0.00001*). No detectable effect of genotype, OT treatment nor experimental paradigm was detected on the number of grooming or rearing episodes. **(e)** In the three-chamber test, when the analysis included both *Oprm1^-/-^* and *Oprm1^+/+^* mice, the time spent in the chamber with the toy was longer in mice trained under the “social” paradigm (*S x P:* *F_1,56_=5.38, p<0.05).*

Results are shown as scatter plots and mean ± sem. Solid stars: significant difference with the vehicle-treated Oprm1^+/+^ group, Tuckey’s post-hoc test following a two-way ANOVA or 2-tailed t-test following a Kruskal-Wallis analysis of variance; open stars: genotype x treatment (Y-maze) or genotype x treatment x stimulus interaction (Social preference - stimulus: mouse/toy or stranger/cage mate comparison), Tukey’s post-hoc test following an analysis of variance (ANOVA); daggers: genotype x treatment interaction; one symbol: p<0.05, two symbols: p<0.01; three symbols: p<0.001. Letters: significant difference with vehicle-treated Oprm1^-/-^ group (2-tailed t-test or Tukey’s post-hoc test); (c): p<0.05, (b): p<0.01, (a): p<0.001. More behavioural parameters in Fig. S4. D: day, M: mouse, T: toy.

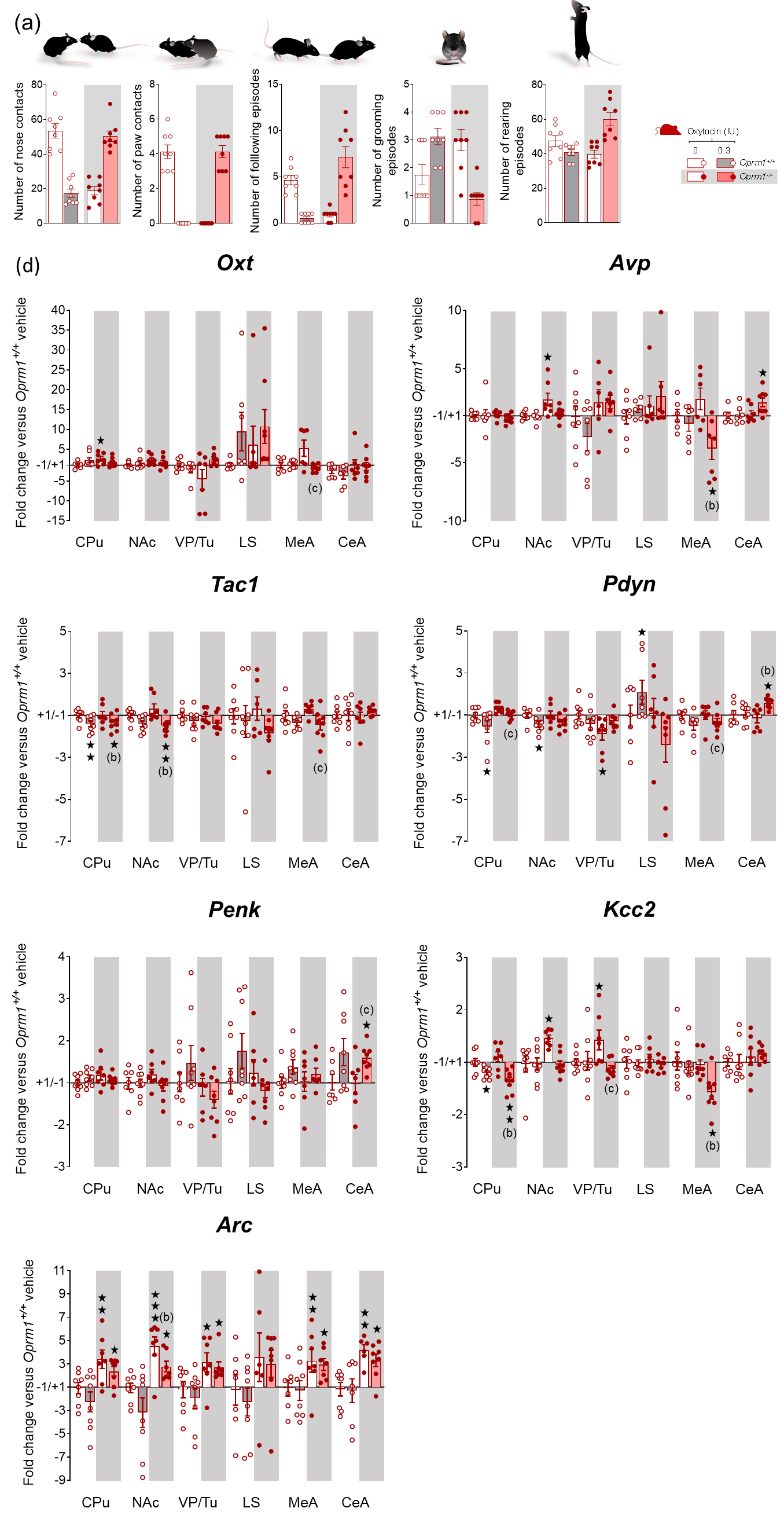

**Figure S5. Transcriptional consequences of social OT conditioning in *Oprm1* null mice and their wild-type controls.** See timeline in Figure 5. (a) OT exposure decreased social interaction in *Oprm1^+/+^* mice while rescuing it in *Oprm1^-/-^* mice, as illustrated by opposite effects on the number of nose contacts (*F_1,28_=119.3, p<0.0001*), paw contacts (*H_3,32_=27.0, p<0.0001)*, following episodes (*H_3,32_=24.9, p<0.0001*) and grooming episodes (*H_3,32_=16.9, p<0.001)*. Per nasal OT increased the number of rearing episodes in *Oprm1^-/-^* mice (*F_1,28_=21.9, p<0.0001)*. (b) Genotype and OT treatment had little impact on the expression of oxytocin and vasopressin genes. Expression of *Kcc2* was increased in the Nac and VP/Tu of *Oprm1* null mice; OT decreased this expression in the CPu of *Oprm1^+/+^* and *Oprm1^-/-^* mice and in the MeA of *Oprm1^-/-^* mice. This treatment reduced *Kcc2* expression in the VP/Tu of *Oprm1^-/-^* mice when compared to vehicle-treated mutant mice. OT treatment reduced *Tac1* expression in the CPu of *Oprm1^+/+^* mice, and in the CPu, NAc and MeA (compared to vehicle-treated *Oprm1* null mice) of *Oprm1^-/-^* mice. OT administration decreased *Pdyn* expression in the CPu and NAc but increased it in the LS of *Oprm1^+/+^* mice; this expression was found decreased in the VP/Tu of vehicle-treated *Oprm1^-/-^* mice and in the MeA of OT-treated *Oprm1^-/-^* mice (compared to vehicle-treated *Oprm1* null mice) but increased in the CeA of OT-treated *Oprm1^-/-^* mice. Genotype and OT treatment had little impact on *Penk* expression. The expression of the immediate early gene *Arc* was found upregulated in the CPu, NAc, VP/Tu, MeA and CeA of *Oprm1^-/-^* mice; OT treatment reduced this expression in the NAc.

Gene expression data are expressed as fold change versus *Oprm1^+/+^* - vehicle group (clustering or scatter plots and mean ± SEM). Comparison to *Oprm1^+/+^* - vehicle group (two-tailed t-test): One star p<0.05, two stars p<0.01, three stars p<0.001. Letters: significant difference with vehicle-treated *Oprm1^-/-^* group (2-tailed t-test); (c): p<0.05, (b): p<0.01, (a): p<0.001. qRT-PCR data are displayed in Table S2.
