## Supplementary material for "Acute, chronic and conditioned effects of intranasal oxytocin in the mu opioid receptor knockout mouse model of autism: social context matters": Table S1

**Table S1. List of primers used for qRT-PCR**

| Refseq | Gene name | Gen title | Forward oligonucleotide | Reverse oligonucleotide |
|---|---|---|---|---|
| NM_009630 | adenosine A2a receptor | *Adora2a* | TCAGCCTCTTGGCTATTGCC | CTCAAACAGACAGGTCACCCG |
| NM_009630 | activity regulated cytoskeletal-associated protein | *Arc* | CCAGGAGAATGACACCAG | TTCAGGAGAAGAGAGGATG |
| NM_009732 | arginine vasopressine | *Avp* | ACACTACGCTCTTCCGCTTGT | CACTGTCTCAGCTCCATGTCA |
| NM_016847 | arginine vasopressine receptor 1A | *Avpr1a* | GGAGAAACGGGAGACAGACA | AAGCCCATTGTACAGCCCAAG |
| NM_011924 | arginine vasopressine receptor 1B | *Avpr1b* | CTGCCTTCAGGTTCTTGCCT | TAATTCACAGGTCATGCGCCA |
| NM_205769 | corticotropin releasing hormone | *Crh* | AGGAGGCATCCTGAGAGAAGT | ATGTTAGGGGCGCTCTCTTC |
| NM_010076 | dopamine receptor D1A | *Drd1a* | AGATCGGGCATTTGGAGAG | GGATGCTGCCTCTTCTTCTG |
| NM_010077 | dopamine receptor D2 | *Drd2* | TGCCATTGTTCTTGGTGTGT | GTGAAGGCGCTGTAGAGGAC |
| NM_010234 | FBJ osteosarcoma oncogene | *Fos* | GAAGGGAACGGAATAAGATG | CATCTTCAAGTTGATCTGTCTC |
| NM_001160353 | glutamate receptor, metabotropic 2 | *Grm2* | CTTGTAGCTATGCCCCGTGT | GACTGGAAGCACCTTTGCAT |
| NM_001013385 | glutamate receptor, metabotropic 4 | *Grm4* | CTTTCTCTGCTATGCCACCACC | TAGCTGATGCTCATGCCAAGCC |
| NM_011025 | oxytocin | *Oxt* | CTGCTTGGCTTACTGGCTCT | GGGAGACACTTGCGCATATC |
| NM_001081147 | oxytocin receptor | *Oxtr* | CTTAGGGCCAAAAGGTGTCA | GCAGGTTTCTATGCCCTCTG |
| NM_018863 | prodynorphin | *Pdyn* | TTTGGCAACGGAAAAGAATC | TAGCGTTTGGCCTGTTTTCT |
| NM_001002927 | preproenkephalin | *Penk* | ATGCAGATGAGGGAGACACC | GCTTCTGCAGCTCTTTTGCT |
| NM_009194 | solute carrier family 12, member 2 | *Slc12a2* | AGGTAAAACATCCGGTGGGT | AGCACAAAGAGAAAGACGCAAC |
| NM_020333 | solute carrier family 12, member 5 | *Slc12a5* | CAGACCTATGTGCAGGGCAA | CCGAGTCGGGATGCGAAATA |
| NM_080853 | solute carrier family 17 (sodium-dependent inorganic phosphate cotransporter), member 6 | *Slc17a6* | AGGCCCTGCTACTGCAAATA | GACACAAAGCAGAGAGGGACT |
| NM_182993 | solute carrier family 17 (sodium-dependent inorganic phosphate cotransporter), member 7 | *Slc17a7* | GGCCATTTGTTGTGTGTCCC | CTCACCCCCACCCAGATTTC |
| NM_009311 | tachykinin 1 | *Tac1* | CCGTTCACTGCTCACTGACACAG | CTCGTTTCCACTCAACTGTTTGC |
