## Supplementary material for "Acute, chronic and conditioned effects of intranasal oxytocin in the mu opioid receptor knockout mouse model of autism: social context matters": Table S2

**Table S2**: Transcription levels of a set of 19 genes in the CPu, NAc, VP/Tu, CeA, MeA, VP and LS in Oprm1+/+ or Oprm1-/- mice treated chronically with OT versus vehicle

| | | | NAc-Oprm1$^{-/-}$/Vehicle | | | | NAc-Oprm1$^{+/+}$/OXT | | | | NAc-Oprm1$^{-/-}$/OXT | | | | CPu-Oprm1$^{-/-}$/Vehicle | | | | CPu-Oprm1$^{+/+}$/OXT | | | | CPu-Oprm1$^{-/-}$/OXT | | | |
|---|---|---|---|---|---|---|---|---|---|---|---|---|---|---|---|---|---|---|---|---|---|---|---|---|---|---|
| Refseq | Gene name | Gen title | FC | sem | p values | Benjamini-Hochberg Adjusted P value | FC | sem | p values | Benjamini-Hochberg Adjusted P value | FC | sem | p values | Benjamini-Hochberg Adjusted P value | FC | sem | p values | Benjamini-Hochberg Adjusted P value | FC | sem | p values | Benjamini-Hochberg Adjusted P value | FC | sem | p values | Benjamini-Hochberg Adjusted P value |
| NM_009630 | adenosine A2a receptor | *Adora2a* | -1.17 | 0.69 | 0.196 | 0.727 | -1.94 | 0.21 | 0.009 | 0.399 | -2.09 | 0.22 | 0.004 | 0.090 | -1.01 | 0.13 | 0.977 | 0.977 | -1.63 | 0.08 | 0.000 | 0.001 | -1.64 | 0.09 | 0.000 | 0.000 |
| NM_009630 | activity regulated cytoskeletal-associated protein | *Arc* | 4.52 | 0.19 | 0.238 | 0.025 | -3.19 | 1.28 | 0.139 | 0.726 | 2.75 | 0.42 | 0.010 | 0.090 | 3.86 | 0.70 | 0.005 | 0.071 | -2.26 | 0.86 | 0.249 | 0.393 | 2.35 | 0.41 | 0.041 | 0.080 |
| NM_009732 | arginine vasopressine | *Avp* | 2.37 | 0.79 | 0.001 | 0.197 | -1.15 | 0.15 | 0.473 | 0.618 | 1.18 | 0.19 | 0.488 | 0.090 | 1.07 | 0.14 | 0.713 | 0.887 | 1.05 | 0.46 | 0.912 | 0.912 | -1.25 | 0.13 | 0.170 | 0.293 |
| NM_016847 | arginine vasopressine receptor 1A | *Avpr1a* | -1.12 | 0.28 | 0.701 | 0.904 | -1.69 | 0.49 | 0.242 | 0.227 | 1.43 | 0.39 | 0.437 | 0.094 | 1.72 | 0.24 | 0.167 | 0.462 | -1.96 | 0.19 | 0.030 | 0.089 | -2.40 | 0.51 | 0.032 | 0.068 |
| NM_011924 | arginine vasopressine receptor 1B | *Avpr1b* | 1.24 | 0.35 | 0.145 | 0.347 | -1.49 | 0.18 | 0.032 | 0.726 | -1.04 | 0.09 | 0.708 | 0.094 | -1.30 | 0.26 | 0.306 | 0.703 | -1.72 | 0.18 | 0.006 | 0.056 | -2.21 | 0.34 | 0.005 | 0.022 |
| NM_205769 | corticotropin releasing hormone | *Crh* | -1.08 | 0.49 | 0.052 | 0.923 | 1.18 | 0.47 | 0.694 | 0.105 | 1.37 | 0.26 | 0.300 | 0.309 | 1.08 | 0.28 | 0.886 | 0.970 | 1.07 | 0.07 | 0.840 | 0.904 | -1.01 | 0.31 | 0.934 | 0.934 |
| NM_010076 | dopamine receptor D1A | *Drd1a* | 1.12 | 0.16 | 0.825 | 0.667 | -1.61 | 0.15 | 0.007 | 0.820 | -1.47 | 0.13 | 0.020 | 0.309 | 1.04 | 0.07 | 0.628 | 0.887 | -1.26 | 0.08 | 0.017 | 0.080 | -1.38 | 0.04 | 0.000 | 0.000 |
| NM_010077 | dopamine receptor D2 | *Drd2* | 1.33 | 0.57 | 0.812 | 0.384 | -1.13 | 0.10 | 0.718 | 0.726 | -1.44 | 0.12 | 0.134 | 0.309 | -1.04 | 0.10 | 0.733 | 0.887 | -1.25 | 0.08 | 0.045 | 0.115 | -1.59 | 0.06 | 0.000 | 0.000 |
| NM_010234 | FBJ osteosarcoma oncogene | *Fos* | 1.49 | 0.09 | 0.105 | 0.421 | -1.25 | 0.21 | 0.465 | 0.820 | -1.01 | 0.24 | 0.917 | 0.309 | 1.51 | 0.24 | 0.144 | 0.462 | -1.60 | 0.28 | 0.110 | 0.211 | -1.27 | 0.13 | 0.243 | 0.350 |
| NM_00116035 | glutamate receptor, metabotropic 2 | *Grm2* | -1.06 | 0.55 | 0.032 | 0.904 | -1.06 | 0.19 | 0.727 | 0.354 | -1.21 | 0.16 | 0.269 | 0.535 | 1.09 | 0.09 | 0.713 | 0.887 | -1.10 | 0.11 | 0.813 | 0.904 | -1.59 | 0.30 | 0.206 | 0.316 |
| NM_00101338 | glutamate receptor, metabotropic 4 | *Grm4* | -1.39 | 0.05 | 0.013 | 0.197 | -1.43 | 0.22 | 0.107 | 0.726 | -1.68 | 0.22 | 0.017 | 0.535 | 1.05 | 0.12 | 0.821 | 0.944 | -1.23 | 0.22 | 0.363 | 0.491 | -1.43 | 0.13 | 0.030 | 0.068 |
| NM_011025 | oxytocin | *Oxt* | 2.25 | 0.06 | 0.080 | 0.241 | 1.59 | 0.56 | 0.411 | 0.721 | 1.64 | 0.54 | 0.360 | 0.535 | 2.61 | 0.55 | 0.024 | 0.142 | 2.24 | 0.61 | 0.107 | 0.211 | 1.72 | 0.39 | 0.178 | 0.293 |
| NM_00108114 | oxytocin receptor | *Oxtr* | 1.04 | 0.16 | 0.712 | 0.904 | -1.32 | 0.28 | 0.333 | 0.227 | -1.24 | 0.17 | 0.314 | 0.535 | 1.21 | 0.14 | 0.481 | 0.790 | -1.28 | 0.16 | 0.235 | 0.393 | -1.66 | 0.20 | 0.028 | 0.068 |
| NM_018863 | prodynorphin | *Pdyn* | 1.19 | 0.11 | 0.034 | 0.472 | -1.00 | 0.1 | 0.989 | 0.820 | -1.07 | 0.12 | 0.743 | 0.535 | 1.18 | 0.12 | 0.180 | 0.462 | 1.11 | 0.06 | 0.256 | 0.393 | 1.07 | 0.04 | 0.394 | 0.453 |
| NM_00100292 | preproenkephalin | *Penk* | 1.00 | 0.32 | 0.888 | 0.923 | -1.43 | 0.14 | 0.049 | 0.989 | -1.34 | 0.13 | 0.107 | 0.535 | 1.33 | 0.08 | 0.044 | 0.206 | -1.53 | 0.28 | 0.110 | 0.211 | 1.01 | 0.07 | 0.932 | 0.934 |
| NM_009194 | solute carrier family 12, member 2 | *Slc12a2* | -1.17 | 0.12 | 0.287 | 0.306 | -1.18 | 0.18 | 0.344 | 0.989 | -1.22 | 0.11 | 0.129 | 0.624 | -1.35 | 0.08 | 0.008 | 0.071 | -1.37 | 0.09 | 0.009 | 0.056 | -1.71 | 0.14 | 0.000 | 0.003 |
| NM_020333 | solute carrier family 12, member 5 | *Slc12a5* | 1.45 | 0.18 | 0.923 | 0.157 | -1.03 | 0.11 | 0.986 | 0.820 | -1.04 | 0.07 | 0.938 | 0.854 | 1.10 | 0.09 | 0.374 | 0.717 | -1.20 | 0.03 | 0.025 | 0.089 | -1.35 | 0.07 | 0.003 | 0.018 |
| NM_080853 | solute carrier family 17 (sodium-dependent inorganic phosphate cotransporter), member 6 | *Slc17a6* | -1.19 | 0.22 | 0.200 | 0.904 | -1.40 | 0.73 | 0.679 | 0.354 | 1.03 | 0.52 | 0.912 | 0.854 | 2.14 | 0.29 | 0.009 | 0.071 | -1.25 | 0.39 | 0.600 | 0.767 | 1.597 | 0.45 | 0.287 | 0.367 |
| NM_182993 | solute carrier family 17 (sodium-dependent inorganic phosphate cotransporter), member 7 | *Slc17a7* | 1.77 | 0.09 | 0.435 | 0.384 | -1.56 | 0.18 | 0.097 | 0.820 | -1.26 | 0.17 | 0.372 | 0.938 | 1.56 | 0.64 | 0.427 | 0.756 | -2.93 | 0.65 | 0.031 | 0.089 | -2.45 | 0.31 | 0.009 | 0.029 |
| NM_009311 | tachykinin 1 | *Tac1* | 1.31 | 0.13 | 0.172 | 0.384 | -1.40 | 0.13 | 0.042 | 0.227 | -1.48 | 0.12 | 0.011 | 0.938 | 1.16 | 0.09 | 0.102 | 0.391 | -1.23 | 0.06 | 0.007 | 0.056 | -1.30 | 0.09 | 0.011 | 0.032 |

| | CeA-Oprm1$^{-/-}$/Vehicle | | | | CeA-Oprm1$^{+/+}$/OXT | | | | CeA-Oprm1$^{-/-}$/OXT | | | | LS-Oprm1$^{-/-}$/Vehicle | | | | LS-Oprm1$^{+/+}$/OXT | | | | LS-Oprm1$^{-/-}$/OXT | | | |
|---|---|---|---|---|---|---|---|---|---|---|---|---|---|---|---|---|---|---|---|---|---|---|---|---|
| Gen title | FC | sem | p values | Benjamini-Hochberg Adjusted P value | FC | sem | p values | Benjamini-Hochberg Adjusted P value | FC | sem | p values | Benjamini-Hochberg Adjusted P value | FC | sem | p values | Benjamini-Hochberg Adjusted P value | FC | sem | p values | Benjamini-Hochberg Adjusted P value | FC | sem | p values | Benjamini-Hochberg Adjusted P value |
| *Adora2a* | -1.28 | 0.15 | 0.503 | 0.925 | -1.51 | 0.31 | 0.296 | 0.858 | 1.05 | 0.17 | 0.824 | 0.824 | -1.36 | 0.75 | 0.641 | 0.827 | 1.44 | 1.37 | 0.904 | 0.951 | -2.21 | 0.84 | 0.290 | 0.574 |
| *Arc* | 4.18 | 0.42 | 0.001 | 0.023 | -1.30 | 0.96 | 0.912 | 0.983 | 3.29 | 0.53 | 0.007 | 0.159 | 3.58 | 1.61 | 0.260 | 0.827 | -2.26 | 1.07 | 0.575 | 0.910 | 3.00 | 1.03 | 0.218 | 0.558 |
| *Avp* | 1.18 | 0.26 | 0.712 | 0.925 | 1.12 | 0.28 | 0.831 | 0.983 | 2.15 | 0.36 | 0.039 | 0.824 | 1.86 | 0.70 | 0.351 | 0.827 | 1.59 | 0.24 | 0.361 | 0.910 | 2.676 | 1.04 | 0.200 | 0.558 |
| *Avpr1a* | 1.08 | 0.16 | 0.865 | 0.925 | -1.15 | 0.29 | 0.601 | 0.983 | 1.12 | 0.17 | 0.775 | 0.418 | 1.62 | 0.25 | 0.222 | 0.827 | 1.22 | 0.24 | 0.626 | 0.910 | 1.26 | 0.10 | 0.494 | 0.617 |
| *Avpr1b* | -1.14 | 0.16 | 0.497 | 0.925 | -1.57 | 0.21 | 0.038 | 0.296 | -1.2 | 0.08 | 0.236 | 0.824 | -1.28 | 0.24 | 0.442 | 0.827 | -1.03 | 0.42 | 0.951 | 0.951 | -2.49 | 0.57 | 0.049 | 0.424 |
| *Crh* | -1.10 | 0.13 | 0.809 | 0.925 | 1.60 | 0.25 | 0.203 | 0.858 | 1.69 | 0.27 | 0.154 | 0.530 | 1.21 | 0.44 | 0.640 | 0.827 | 2.03 | 0.48 | 0.091 | 0.770 | 1.66 | 0.63 | 0.359 | 0.574 |
| *Drd1a* | 1.05 | 0.13 | 0.859 | 0.925 | -1.21 | 0.20 | 0.511 | 0.983 | 1.23 | 0.05 | 0.368 | 0.748 | 1.69 | 0.42 | 0.625 | 0.827 | 2.05 | 0.33 | 0.196 | 0.770 | -1.02 | 0.31 | 0.350 | 0.574 |
| *Drd2* | -1.38 | 0.26 | 0.225 | 0.925 | 1.03 | 0.33 | 0.983 | 0.983 | 1.13 | 0.13 | 0.650 | 0.414 | -1.36 | 0.12 | 0.091 | 0.827 | 1.10 | 0.13 | 0.598 | 0.910 | -1.19 | 0.15 | 0.399 | 0.574 |
| *Fos* | 1.47 | 0.23 | 0.081 | 0.624 | 1.19 | 0.18 | 0.373 | 0.858 | 1.20 | 0.11 | 0.216 | 0.685 | 1.22 | 0.61 | 0.802 | 0.827 | 1.65 | 1.04 | 0.628 | 0.910 | -1.97 | 0.51 | 0.332 | 0.574 |
| *Grm2* | -1.54 | 0.20 | 0.038 | 0.446 | -1.24 | 0.16 | 0.259 | 0.858 | 1.14 | 0.18 | 0.547 | 0.450 | 1.23 | 0.09 | 0.152 | 0.898 | -1.04 | 0.16 | 0.833 | 0.951 | 1.15 | 0.03 | 0.136 | 0.558 |
| *Grm4* | -1.46 | 0.24 | 0.125 | 0.720 | 1.03 | 0.41 | 0.933 | 0.983 | 1.28 | 0.20 | 0.283 | 0.685 | 1.07 | 0.07 | 0.551 | 0.827 | 1.22 | 0.13 | 0.225 | 0.770 | -1.03 | 0.08 | 0.862 | 0.862 |
| *Oxt* | -1.02 | 0.42 | 0.931 | 0.931 | -1.70 | 0.47 | 0.298 | 0.858 | 1.51 | 0.63 | 0.566 | 0.340 | 6.16 | 3.87 | 0.295 | 0.827 | 9.46 | 4.08 | 0.113 | 0.770 | 10.75 | 3.80 | 0.055 | 0.424 |
| *Oxtr* | -1.33 | 0.33 | 0.322 | 0.925 | -1.44 | 0.15 | 0.031 | 0.296 | -1.20 | 0.10 | 0.135 | 0.110 | 1.14 | 0.06 | 0.393 | 0.827 | -1.02 | 0.10 | 0.927 | 0.951 | 1.18 | 0.05 | 0.252 | 0.574 |
| *Pdyn* | -1.03 | 0.23 | 0.866 | 0.925 | 1.72 | 0.32 | 0.092 | 0.530 | 1.59 | 0.10 | 0.016 | 0.340 | 1.24 | 0.26 | 0.647 | 0.827 | 1.76 | 0.36 | 0.178 | 0.770 | -1.19 | 0.14 | 0.509 | 0.617 |
| *Penk* | -1.15 | 0.20 | 0.533 | 0.925 | 1.06 | 0.16 | 0.879 | 0.983 | 1.58 | 0.09 | 0.019 | 0.110 | 1.10 | 0.56 | 0.859 | 0.898 | 2.08 | 0.47 | 0.170 | 0.770 | -2.41 | 0.73 | 0.200 | 0.558 |
| *Slc12a2* | -1.04 | 0.18 | 0.805 | 0.925 | 1.06 | 0.19 | 0.831 | 0.983 | 1.18 | 0.21 | 0.528 | 0.685 | 1.00 | 0.11 | 0.978 | 0.827 | -1.15 | 0.24 | 0.633 | 0.910 | -1.22 | 0.15 | 0.397 | 0.574 |
| *Slc12a5* | 1.10 | 0.15 | 0.548 | 0.925 | 1.00 | 0.14 | 0.966 | 0.983 | 1.16 | 0.05 | 0.082 | 0.243 | 1.05 | 0.07 | 0.683 | 0.827 | -1.02 | 0.06 | 0.905 | 0.951 | -1.04 | 0.04 | 0.779 | 0.814 |
| *Slc17a6* | 1.14 | 0.31 | 0.822 | 0.925 | 1.01 | 0.10 | 0.731 | 0.983 | -1.76 | 0.29 | 0.017 | 0.110 | 1.08 | 0.34 | 0.853 | 0.978 | 1.20 | 0.22 | 0.539 | 0.910 | 1.27 | 0.30 | 0.504 | 0.617 |
| *Slc17a7* | -1.08 | 0.29 | 0.754 | 0.925 | -1.60 | 0.22 | 0.020 | 0.296 | -1.30 | 0.15 | 0.084 | 0.243 | -1.18 | 0.21 | 0.620 | 0.827 | -1.21 | 0.18 | 0.522 | 0.910 | -1.19 | 0.43 | 0.761 | 0.814 |
| *Tac1* | -1.10 | 0.24 | 0.706 | 0.925 | 1.03 | 0.27 | 0.948 | 0.983 | 1.20 | 0.06 | 0.293 | 0.450 | 1.32 | 0.43 | 0.574 | 0.898 | -1.29 | 0.67 | 0.774 | 0.951 | -1.85 | 0.27 | 0.107 | 0.558 |

| | MeA-Oprm1$^{-/-}$/Vehicle | | | | MeA-Oprm1$^{+/+}$/OXT | | | | MeA-Oprm1$^{-/-}$/OXT | | | | VP-Oprm1$^{-/-}$/Vehicle | | | | VP-Oprm1$^{+/+}$/OXT | | | | VP-Oprm1$^{-/-}$/OXT | | | |
|---|---|---|---|---|---|---|---|---|---|---|---|---|---|---|---|---|---|---|---|---|---|---|---|---|
| Gen title | FC | sem | p values | Benjamini-Hochberg Adjusted P value | FC | sem | p values | Benjamini-Hochberg Adjusted P value | FC | sem | p values | Benjamini-Hochberg Adjusted P value | FC | sem | p values | Benjamini-Hochberg Adjusted P value | FC | sem | p values | Benjamini-Hochberg Adjusted P value | FC | sem | p values | Benjamini-Hochberg Adjusted P value |
| *Adora2a* | -1.13 | 0.15 | 0.642 | 0.896 | -1.66 | 0.10 | 0.045 | 0.259 | -2.10 | 0.36 | 0.028 | 0.066 | -3.00 | 0.72 | 0.039 | 0.168 | -2.42 | 0.98 | 0.221 | 0.648 | -6.34 | 1.21 | 0.001 | 0.006 |
| *Arc* | 4.08 | 0.64 | 0.008 | 0.093 | -1.16 | 0.87 | 0.938 | 0.938 | 2.93 | 0.47 | 0.034 | 0.072 | 3.14 | 0.81 | 0.043 | 0.168 | -1.86 | 0.91 | 0.573 | 0.776 | 2.69 | 0.45 | 0.044 | 0.147 |
| *Avp* | 2.44 | 0.85 | 0.218 | 0.750 | -1.67 | 0.58 | 0.409 | 0.587 | -3.80 | 0.86 | 0.027 | 0.066 | 2.14 | 1.00 | 0.401 | 0.710 | -2.81 | 1.08 | 0.248 | 0.648 | 2.19 | 0.53 | 0.271 | 0.389 |
| *Avpr1a* | 1.12 | 0.16 | 0.540 | 0.887 | -1.77 | 0.11 | 0.000 | 0.007 | -1.88 | 0.14 | 0.000 | 0.006 | 1.65 | 0.24 | 0.015 | 0.159 | -1.73 | 0.30 | 0.032 | 0.648 | -1.57 | 0.12 | 0.000 | 0.006 |
| *Avpr1b* | -1.21 | 0.16 | 0.437 | 0.839 | -1.59 | 0.17 | 0.041 | 0.259 | -1.36 | 0.18 | 0.204 | 0.247 | 1.05 | 0.11 | 0.821 | 0.908 | -1.74 | 0.37 | 0.085 | 0.648 | -1.05 | 0.19 | 0.842 | 0.881 |
| *Crh* | -1.38 | 0.29 | 0.359 | 0.752 | 1.03 | 0.16 | 0.800 | 0.836 | -1.48 | 0.18 | 0.147 | 0.212 | -1.43 | 0.18 | 0.329 | 0.688 | 1.12 | 0.14 | 0.985 | 0.985 | -1.32 | 0.15 | 0.383 | 0.464 |
| *Drd1a* | -1.41 | 0.5 | 0.539 | 0.887 | -1.61 | 0.25 | 0.201 | 0.385 | -1.69 | 0.19 | 0.118 | 0.182 | -1.51 | 0.25 | 0.225 | 0.552 | -1.48 | 0.29 | 0.282 | 0.648 | -2.62 | 0.37 | 0.005 | 0.020 |
| *Drd2* | -1.35 | 0.3 | 0.293 | 0.750 | -1.30 | 0.16 | 0.149 | 0.358 | -2.22 | 0.26 | 0.001 | 0.009 | -1.02 | 0.15 | 0.864 | 0.908 | -1.16 | 0.17 | 0.469 | 0.756 | -2.04 | 0.21 | 0.001 | 0.006 |
| *Fos* | -1.05 | 0.24 | 0.803 | 0.939 | -1.21 | 0.19 | 0.453 | 0.614 | -1.35 | 0.26 | 0.316 | 0.339 | -1.04 | 0.21 | 0.868 | 0.908 | -1.13 | 0.14 | 0.493 | 0.756 | -1.19 | 0.03 | 0.201 | 0.331 |
| *Grm2* | -1.39 | 0.27 | 0.284 | 0.750 | -1.18 | 0.16 | 0.559 | 0.676 | -1.44 | 0.13 | 0.095 | 0.156 | 1.27 | 0.21 | 0.240 | 0.552 | 1.07 | 0.27 | 0.817 | 0.940 | -1.05 | 0.04 | 0.452 | 0.520 |
| *Grm4* | -1.63 | 0.12 | 0.038 | 0.294 | -1.56 | 0.26 | 0.138 | 0.358 | -2.38 | 0.29 | 0.003 | 0.015 | 1.33 | 0.33 | 0.383 | 0.710 | 1.46 | 0.35 | 0.265 | 0.648 | -1.26 | 0.15 | 0.183 | 0.331 |
| *Oxt* | 5.26 | 1.89 | 0.078 | 0.451 | 1.22 | 0.34 | 0.790 | 0.836 | -1.70 | 0.33 | 0.345 | 0.345 | -4.62 | 2.49 | 0.146 | 0.420 | -1.95 | 0.81 | 0.311 | 0.650 | 2.43 | 0.58 | 0.096 | 0.247 |
| *Oxtr* | 1.16 | 0.25 | 0.596 | 0.896 | -1.49 | 0.26 | 0.124 | 0.358 | -1.85 | 0.21 | 0.005 | 0.021 | 1.03 | 0.17 | 0.934 | 0.934 | -1.48 | 0.25 | 0.109 | 0.648 | -1.86 | 0.17 | 0.001 | 0.006 |
| *Pdyn* | 1.03 | 0.23 | 0.921 | 0.963 | 1.38 | 0.17 | 0.107 | 0.358 | 1.20 | 0.13 | 0.325 | 0.339 | -1.11 | 0.21 | 0.706 | 0.903 | 1.47 | 0.41 | 0.341 | 0.654 | -1.40 | 0.20 | 0.188 | 0.331 |
| *Penk* | -1.00 | 0.20 | 0.98 | 0.980 | -1.50 | 0.22 | 0.089 | 0.358 | -1.34 | 0.18 | 0.199 | 0.247 | -1.89 | 0.29 | 0.020 | 0.159 | -1.34 | 0.25 | 0.270 | 0.648 | -1.29 | 0.14 | 0.245 | 0.376 |
| *Slc12a2* | -1.36 | 0.21 | 0.195 | 0.750 | -1.25 | 0.16 | 0.302 | 0.464 | -1.81 | 0.13 | 0.001 | 0.010 | 1.05 | 0.17 | 0.808 | 0.908 | -1.23 | 0.19 | 0.258 | 0.648 | -1.45 | 0.06 | 0.000 | 0.003 |
| *Slc12a5* | -1.05 | 0.10 | 0.722 | 0.923 | -1.11 | 0.13 | 0.543 | 0.676 | -1.57 | 0.12 | 0.015 | 0.050 | 1.43 | 0.18 | 0.039 | 0.168 | 1.03 | 0.17 | 0.882 | 0.946 | -1.12 | 0.05 | 0.151 | 0.331 |
| *Slc17a6* | 1.01 | 0.13 | 0.857 | 0.939 | -1.57 | 0.24 | 0.156 | 0.358 | -1.58 | 0.10 | 0.054 | 0.104 | -1.1 | 0.19 | 0.705 | 0.903 | -1.11 | 0.17 | 0.634 | 0.811 | -1.04 | 0.10 | 0.841 | 0.881 |
| *Slc17a7* | -1.17 | 0.45 | 0.835 | 0.939 | -1.60 | 0.32 | 0.256 | 0.454 | -2.42 | 0.43 | 0.025 | 0.066 | -6.00 | 1.01 | 0.003 | 0.076 | -1.77 | 1.89 | 0.786 | 0.940 | -2.58 | 1.08 | 0.316 | 0.428 |
| *Tac1* | 1.28 | 0.12 | 0.337 | 0.752 | -1.35 | 0.12 | 0.171 | 0.358 | -1.44 | 0.25 | 0.179 | 0.243 | -1.08 | 0.14 | 0.651 | 0.903 | -1.28 | 0.19 | 0.236 | 0.648 | -1.38 | 0.14 | 0.064 | 0.186 |
